# Predictive coding across the left fronto-temporal hierarchy during language comprehension

**DOI:** 10.1101/2021.02.17.431452

**Authors:** Lin Wang, Lotte Schoot, Trevor Brothers, Edward Alexander, Lena Warnke, Minjae Kim, Sheraz Khan, Matti Hämäläinen, Gina R. Kuperberg

## Abstract

We used MEG and ERPs to track the time-course and localization of evoked activity produced by *expected*, *unexpected plausible* and *implausible* words during incremental language comprehension. We interpret the full pattern of evoked responses and functional connectivity within a hierarchical predictive coding framework in which evoked activity reflects residual information (error) that is passed up and down the cortical hierarchy. Between 300-500ms, the three conditions produced progressively larger responses within left temporal cortex (lexico-semantic prediction error), while *implausible* inputs produced a selectively enhanced response within left inferior frontal cortex (event-level prediction error). Between 600-1000ms, *unexpected plausible* words activated left inferior frontal and middle temporal cortices (feedback activity that induced top-down shifts of event-level and lexico-semantic representations), while highly *implausible* inputs activated posterior fusiform and medial temporal cortices, supporting orthographic reanalysis and new learning. Therefore, predictive coding may provide a unifying theory that links language comprehension to other domains of cognition.

## Introduction

One of the most amazing feats of human cognition is the ability to comprehend language— to transform a stream of linguistic inputs into a high-level representation of real-world events (an *event model*^1^). Language comprehension can be understood as *probabilistic inference*: the use of prior knowledge, encoded within an internal generative model, to infer the underlying representation that best “explains” the bottom-up input^2^. According to an influential theory of brain function, inference is approximated by an optimization algorithm known as *predictive coding*^3–6^.

Predictive coding claims that higher levels of the cortical hierarchy, representing information over longer spatiotemporal scales, continually generate top-down predictions of information at lower cortical levels, which represent information over shorter spatiotemporal scales. As new bottom-up information becomes available to lower levels, the brain computes *prediction error*—residual information within the input that cannot be explained by top-down predictions. This prediction error is then passed back up to update higher-level representations, allowing them to produce more accurate predictions. Inference is complete when, across multiple iterations of the algorithm, prediction error is minimized across all levels of the cortical hierarchy.

In the language domain, studies of speech perception^7, 8^ and visual word recognition^9, 10^ have shown that predictable inputs produce less activity than unpredictable inputs within regions that encode low-level phonological and orthographic representations. This has been taken as evidence for the suppression of low-level prediction error by accurate top-down predictions. An important open question is whether the principles of predictive coding can also account for the spatiotemporal dynamics of activity produced during higher-level language comprehension, which requires us to infer information at *multiple* time-scales as new linguistic inputs unfold in real time. Previous studies have shown that the lexico-semantic features of individual words are inferred within lower levels of the fronto-temporal hierarchy––the left temporal cortex^11^, while higher-level event representations are incrementally updated over a longer time span at higher levels of the hierarchy––the left inferior frontal cortex (IFC)^12^. However, it remains unclear *when* and *how* these lower and higher-level regions are modulated as new information flows up and down the fronto-temporal hierarchy during real-time comprehension.

### 300-500ms

Important clues about the *time-course* of online language comprehension come from event-related potentials (ERPs). Previous ERP studies have consistently shown that the amplitude of the N400^13, 14^––a phase-locked neural response observed at the scalp surface between 300-500ms––is highly sensitive to the lexico-semantic predictability of incoming words; that is, to the degree of match between their semantic properties and those activated by the prior context^13–16^. The N400 is therefore generally smallest to words that are predictable in context, larger to words that are unpredictable but plausible^13, 17^, and largest to implausible words whose semantic features are highly unpredictable^18–20^. However, the neuroanatomical sources of these effects remain unclear. In fMRI, the sluggish hemodynamic response is largely insensitive to transient, feedforward neural activity^21^, like that reflected in the N400^11, 22^. And, although previous intracranial^23^ and MEG studies^24, 25^ show that activity within the 300-500ms (N400) time-window can localize to both temporal and inferior prefrontal cortices, none of these studies have experimentally distinguished between the effects of lexical predictability in plausible sentences, and contextual implausibility. Addressing this question is important because it can help distinguish predictive coding from other theoretical and neurobiological frameworks of language comprehension.

According to a *prediction-integration* account, top-down lexico-semantic predictions facilitate the retrieval of expected incoming words in plausible sentences, while the effects of implausibility on the N400 are attributed to the difficulty in “integrating” these words into their broader context^19, 20^. This functional distinction between lexical prediction and integration would predict a reduction in evoked activity to *expected* words within regions of left temporal cortex that support lexico-semantic processing^11^ (*Expected* < *Unexpected plausible* = *Implausible*), and an enhanced response to *implausible* words within left-IFC, which is thought to mediate higher-level integration processes^12^ (*Implausible > Unexpected plausible* = *Expected*).

In contrast, a *fully predictive* architecture posits that *all* contextual effects are driven by top-down lexical facilitation^26, 27^. This would therefore predict a progressive increase in lexico-semantic activity within left temporal cortex to *expected*, *unexpected plausible* and *implausible* words, but no differences within left-IFC. Finally, others have argued that the N400 reflects incremental updates of a single state^28^, which may be maintained by continuous cycles of feedforward/feedback activity between left temporal and inferior frontal cortices^29^. This *single-state* account would therefore predict progressive increases in evoked activity to *expected*, *unexpected plausible* and *implausible* words within *both* temporal and inferior frontal regions between 300-500ms.

In all these accounts, a basic assumption is that the magnitude of the N400 *directly* indexes the change in state induced by new bottom-up inputs at a given level of representation—a lexico-semantic state^30^, the landscape of semantic memory^14^, or the unfolding event model^28^. Predictive coding makes a different set of assumptions. Like fully predictive architectures^26, 27^, it assumes a top-down flow of information from higher to lower levels of the fronto-temporal hierarchy. Like single-state models^28, 29^, it assumes a continuous feedforward flow of information across this network within the N400 time-window. However, unlike either of these previous frameworks, predictive coding does not attribute the evoked neural response directly to the process of updating these internal states. Instead, it assumes that the phase-locked neural response indexes the magnitude of *prediction error*––the residual information encoded within the newly-updated state that is not present in top-down predictions^5^. This error is explicitly computed in the service of probabilistic inference^3–6^.

Within this predictive coding framework, evoked activity produced between 300-500ms within left temporal cortex indexes *lexico-semantic prediction error*, while activity produced within left-IFC indexes higher-level *event prediction error* (see Figure 1). Like fully predictive and single-state architectures, this framework predicts that, in the N400 time-window, regions of left temporal cortex supporting lexico-semantic processing should show graded increases in activity across *expected*, *unexpected plausible* and *implausible* inputs. This is because the magnitude of lexico-semantic prediction error should become progressively larger due to progressively less suppression by top-down predictions from left-IFC. Unlike these previous frameworks, however, predictive coding predicts that an enhanced evoked response within left-IFC (reflecting event-level prediction error) should only be produced by *implausible* inputs, whose statistics differs strongly from the statistics of natural environmental inputs^4^; that is, when a newly-inferred implausible event cannot be explained/suppressed by top-down predictions based on stored real-world knowledge.

**Figure 1.**
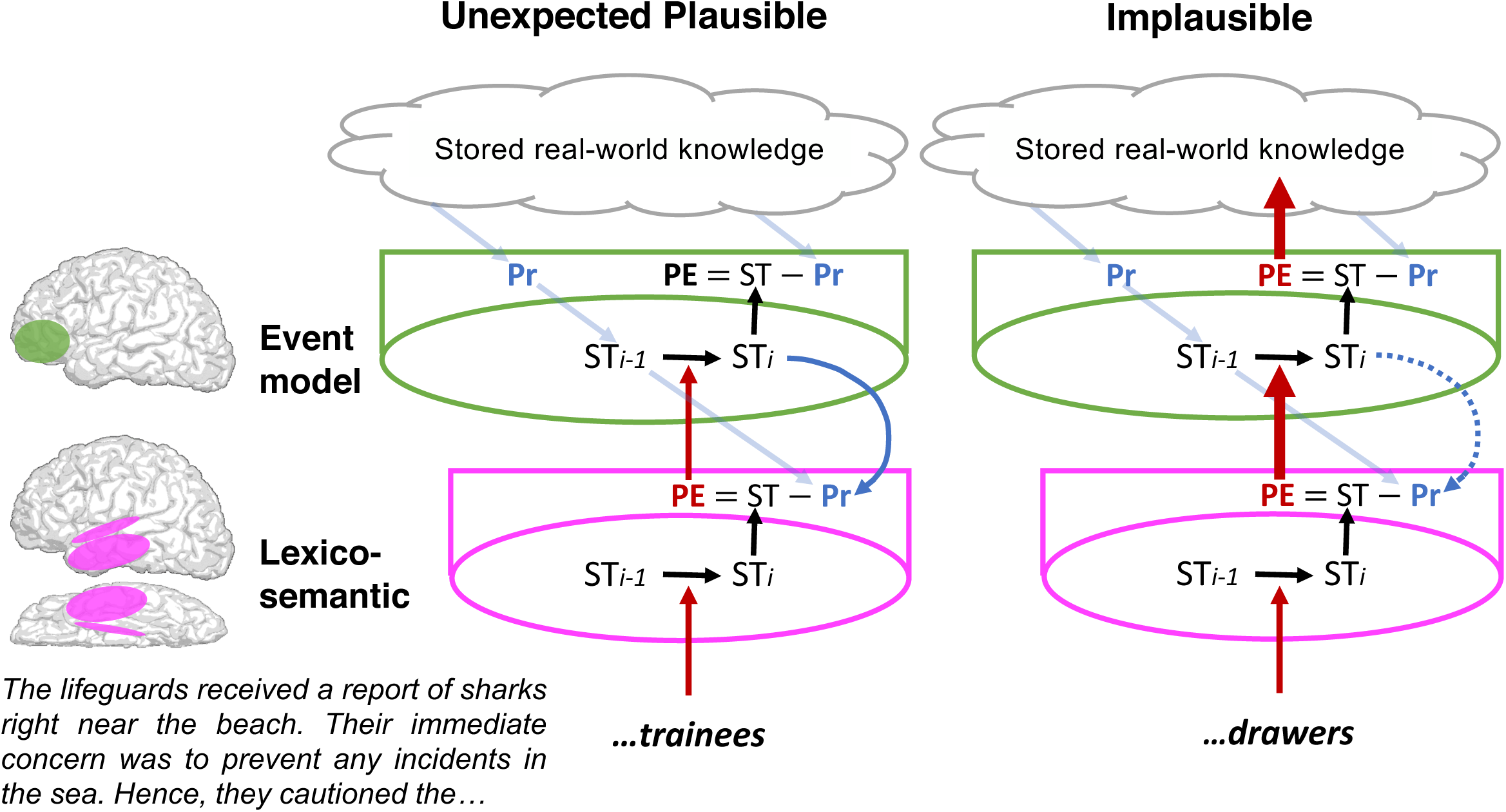
300-500ms (N400): Predictive coding across the left fronto-temporal hierarchy. In constraining contexts, top-down predictions, based on stored real-world and linguistic knowledge, are propagated down the cortical hierarchy (faint blue diagonal arrows) before new bottom-up input becomes available. **Lexico-semantic level** (*pink*): Within left temporal cortex, the lexico-semantic representation of the incoming word is inferred throughout the N400 time-window (ST*_i-1_*→ST*_i_*). Residual lexico-semantic information that cannot be explained by the top-down predictions (PE = max(0, ST– Pr), simplified in the figure as PE = ST – Pr) produces lexico-semantic prediction error, which manifests as an evoked response. Lexico-semantic prediction error, and evoked activity within left temporal cortex is therefore smallest to *expected* words (e.g. “swimmers”, not shown), larger to *unexpected plausible* words (e.g. “trainees”, *left*), where some semantic features (e.g. <animate>) were predicted, and largest to *implausible* words (e.g. “drawers”, *right*) where no semantic features were predicted. **Event level** (*green*): Throughout the N400 time-window, lexico-semantic prediction error passes up to left inferior frontal cortex (IFC) (red arrows), where it induces updates of the event model. *<u>Unexpected plausible</u>* (*left*): These updates yield a plausible interpretation (e.g. <lifeguards cautioned trainees>) that is congruent with predictions about the possibility/plausibility of real-world events. Therefore, higher-level *event prediction error* is suppressed, and the evoked response within left-IFC is no larger than to *expected* inputs. In addition, the newly-updated event model (ST*_i_*) now generates correct top-down predictions (blue curly arrow) that “switch off”/suppress lower-level lexico-semantic prediction error, leading to the reduction of the evoked response within left temporal cortex at the end of the N400 time-window. *<u>Implausible</u>* (*right*): Updates of the event model yield an *implausible* interpretation (e.g. <lifeguards cautioned drawers>) that cannot be explained by real-world knowledge predictions, resulting in a large *event prediction error* and evoked response within left-IFC (red arrow). Moreover, because it is more difficult to converge on an implausible interpretation, new top-down lexico-semantic predictions are less likely to suppress lower-level lexico-semantic prediction error (blue dotted curly arrow), further enhancing and prolonging the evoked response within left temporal cortex.

### 600-1000ms

Previous ERP studies have also shown that unpredictable inputs can produce additional evoked responses between 600-1000ms, particularly in rich, schema-constraining contexts. This later-stage activity manifests at the scalp surface as a set of positive-going waveforms between 600-1000ms, with different distributions depending on whether the interpretation is plausible or implausible^18, 31–33^. Unexpected words that yield plausible interpretations produce a late frontally-distributed positivity^18, 33, 34^, while anomalous inputs that yield highly implausible interpretations produce a late posterior positivity, otherwise known as the P600^18, 35, 36^.

It has been proposed that the *late frontal positivity* reflects lexical inhibition^34^, or high-level shifts of the event model^18, 33^, while the *late posterior positivity/P600* reflects a reanalysis of the perceptual input that is triggered by linguistic conflict^18, 33, 35, 36^. However, we know very little about the neuroanatomical sources of this later activity. Most previous MEG and intracranial studies have failed to report activity beyond the N400 time-window. And although fMRI may be more sensitive to later-stage feedback activity than earlier feedforward transient responses (like the N400)^21^, the sluggish hemodynamic response additionally captures still later activity that extends well beyond 1000ms, indexing the processing of subsequent words and offline reflections about the meaning of sentences as a whole.

Hierarchical predictive coding provides a biologically-motivated computational framework for understanding this later-stage neural activity. Within a dynamic generative framework, inferring an event that is plausible but inconsistent with prior high-confidence beliefs, leads the brain to infer a *systematic change*, resulting in the retrieval of new schema-relevant information from long-term memory^18, 37^. Predictive coding posits that this will result in the generation of new top-down predictions that are passed down from higher to lower levels of the cortical hierarchy, driving a shift of the event model and lexico-semantic state (see Figure 2, left). This account therefore predicts that *unexpected plausible* continuations will produce late evoked effects within both left inferior frontal and temporal cortices.

**Figure 2.**
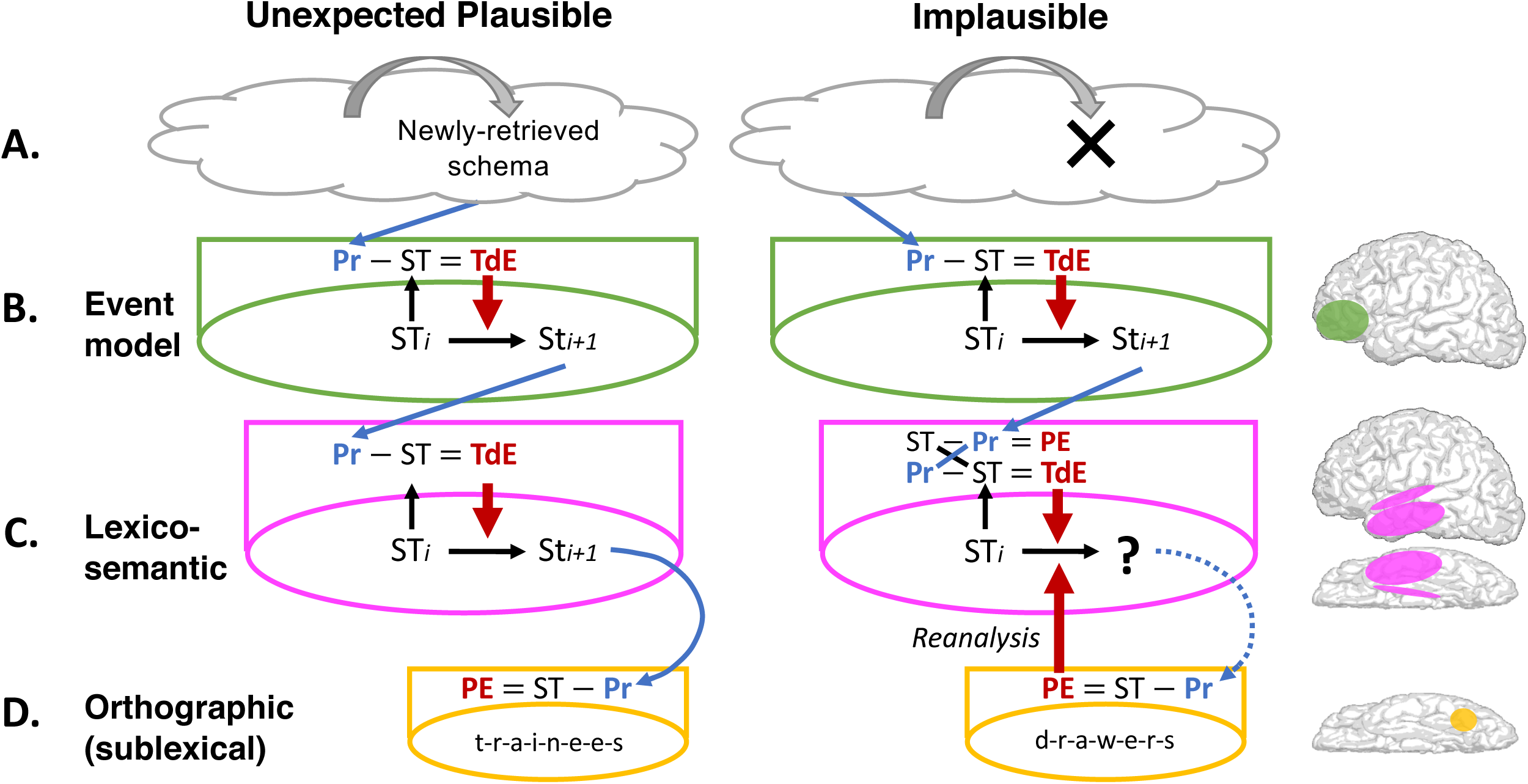
600-1000ms: Predictive coding across the left fronto-temporal hierarchy. **Unexpected plausible (*left panel*).** **A.** The plausible event inferred between 300-500ms (e.g. <lifeguards cautioned trainees>) is out of keeping with the schema previously inferred from the discourse context. This triggers the retrieval of new schema from long-term memory between 600-1000ms (gray curly arrow), which generate new predictions (blue diagonal arrow). **B.** *Event level:* In left-IFC, residual information within the new predictions that cannot be explained by the prior event model produces *top-down event error* (max(0, Pr – ST) = TdE, simplified in the figure as Pr – ST = TdE), which manifests as a late evoked response. This induces a top-down shift of the event model (ST*_i_*→ST*_i+1_*), which generates new predictions (blue diagonal arrow). **C.** *Lexico-semantic level:* In left temporal cortex, residual information within the new predictions that cannot be explained by the prior lexico-semantic state produces *top-down lexico-semantic error*, which manifests as a late evoked response, and induces a top-down lexico-semantic shift. This produces orthographic predictions (blue curly arrow) that continue to suppress orthographic prediction error at the level below (**D**). **Implausible** (*right panel*). **A.** There are no stored schemas within memory that can explain the implausible/impossible input (black cross). Therefore, between 600-1000ms, top-down predictions continue to be generated based on the prior context (blue diagonal arrow). **B.** *Event level:* In left-IFC, event predictions (e.g. <lifeguard cautioned swimmers>) fail to match the implausible event inferred in the N400 time-window (<lifeguards cautioned drawers>), producing *top-down event error* and a late evoked response. This induces a top-down shift to a plausible event (<lifeguard cautioned swimmers>), which generates new lexico-semantic predictions. **C.** *Lexico-semantic level:* In left temporal cortex, these predictions (“swimmers”, <animate>) are incompatible with the lexico-semantic state inferred in the N400 time-window (“drawers”, <inanimate>), resulting in a destabilization of this state (indicated with a “?”), and the production of inaccurate orthographic predictions that are passed down to the level below (blue curly dotted arrow). **D.** *Orthographic level:* Within left posterior fusiform cortex, the inaccurate orthographic predictions fail to suppress prediction error produced by the orthographic state (d-r-a-w-e-r-s), resulting in an evoked response (reanalysis).

However, if the comprehender encounters a highly *implausible* input that yields an anomalous interpretation (e.g. “..*.they cautioned the *drawers*”), the brain will be unable to retrieve new schema. This will result in a failure to switch off prediction error at still lower levels of the cortical hierarchy that encode lower-level linguistic representations. This therefore predicts that highly implausible words will produce later activity within lower-level regions such as the posterior fusiform cortex, which supports sublexical orthographic processing^9, 10^, i.e. orthographic reanalysis.

### This study

We used MEG and ERPs to track the time-course and spatial localization of evoked, phase-locked neural activity produced during the first 1000ms following word onset. Participants read multi-sentence discourse contexts, e.g. *“The lifeguards received a report of sharks right near the beach. Their immediate concern was to prevent any incidents in the sea. Hence, they cautioned the…”.* Critical words in the final sentence were (a) *expected* (e.g., *“swimmers”*), (b) *unexpected* but *plausible* (e.g., *“trainees”*), or (c) highly *implausible* (e.g., *“drawers”*).

To spatiotemporally localize the activity produced by these critical words, we carried out a distributed source localization analysis of the MEG data, which is relatively undistorted by conductivities of the skull and scalp. This enabled us to determine the full time-course of evoked activity produced as information flowed up and down the cortical hierarchy. To test our hypotheses, we compared evoked activity across the three conditions, and also examined the functional connectivity between left temporal and inferior frontal cortices, within the 300-500ms and 600-1000ms time-windows. In addition, we simultaneously collected ERP data. This was important because, although MEG and ERP both measure phase-locked evoked activity, they do not capture precisely the same underlying signal^38^ (see Supplementary Materials). Therefore, by collecting ERP data using the same stimuli and participants, we were able to replicate previous ERP findings^18^, and link them directly to the source-localized effects detected by MEG.

## Results

### Behavioral

Participants correctly judged the plausibility of 89.42% scenarios (SD: 8.83%) and answered 80.12% (SD: 11.50%) of the comprehension questions correctly, suggesting that they were engaged in comprehension (see Supplementary Materials).

### ERP

The ERP results (Figure 3, Table 1) replicate previous findings^18^. Between 300-500ms, the N400 amplitude increased across the three conditions. Between 600-1000ms, the *unexpected plausible* words produced a larger *late frontal positivity* than both the *expected* and *implausible* continuations, while the *implausible* words produced a larger *late posterior positivity/P600* than both the *expected* and *unexpected plausible* continuations.

**Figure 3.**
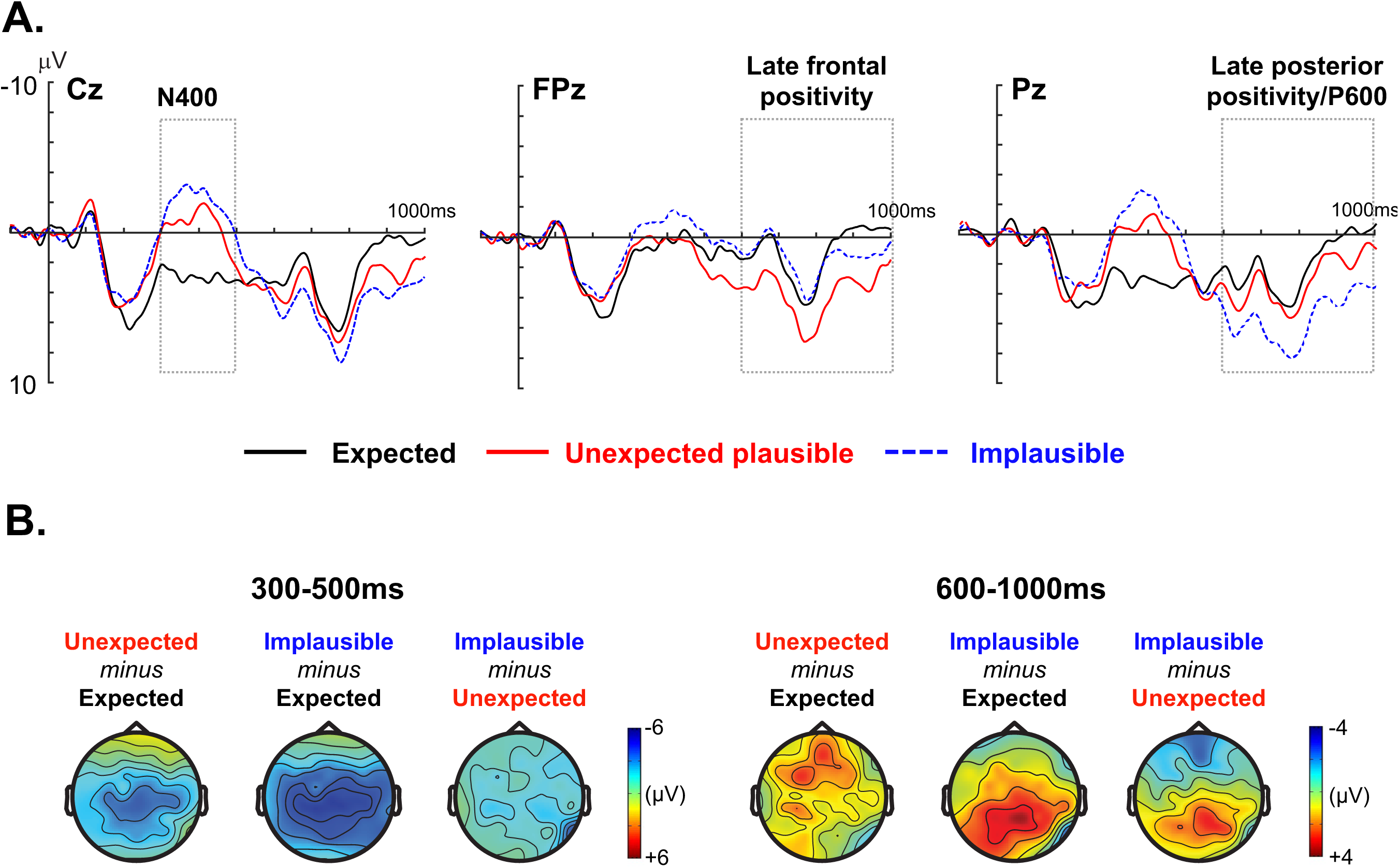
ERP results. **A. Grand-averaged ERP waveforms** elicited by critical words in each of the three conditions, shown at three representative electrode sites: Cz, FPz and Pz. *Expected*: solid black line; *Unexpected plausible*: solid red line; *Implausible*: dashed blue line. Negative voltage is plotted upwards. Dotted boxes are used to indicate the time-windows corresponding to the N400 (300-500ms), the *late frontal positivity* (600-1000ms) and the *late posterior positivity/P600* (600-1000ms) ERP components. **B. Voltage maps** show the topographic distributions of the ERP effects produced by contrasting *Expected*, *Unexpected plausible* and *Implausible* critical words between 300-500ms (left panel) and between 600-1000ms (right panel). Note that the N400 effects and the late positivity effects are shown at different voltage scales to better illustrate the scalp distribution of each effect.

**Table 1.** ERP statistical results.

| **Spatiotemporal Region** | **Contrast** | ***t*-value**<br>***df*(31)** | ***p*-value** | **Effect size (*d*)** |
| --- | --- | --- | --- | --- |
| N400<br>(Central region,<br>300-500ms) | Unexpected > Expected | -5.31 | < 0.001 | 0.94 |
|  | Implausible > Expected | -7.72 | < 0.001 | 1.36 |
|  | Implausible > Unexpected | -4.63 | 0.001 | 0.82 |
| *Late frontal positivity*<br>(Prefrontal region,<br>600-1000ms) | Unexpected > Expected | 3.03 | 0.005 | 0.53 |
|  | Implausible = Expected | 1.51 | 0.14 | 0.24 |
|  | Unexpected > Implausible | 2.52 | 0.018 | 0.42 |
| *Late posterior positivity/P600*<br>(Posterior region,<br>600-1000ms) | Unexpected = Expected | 1.91 | 0.07 | 0.34 |
|  | Implausible > Expected | 7.65 | < 0.001 | 1.12 |
|  | Implausible > Unexpected | 5.75 | < 0.001 | 0.99 |

### MEG

Between 300-500ms, the sensor-level MEG findings showed a similar graded increase in evoked activity, see Figure 4A (the evoked response to the *implausible* continuations was larger in MEG than in ERP, see Supplementary Materials for discussion). Between 600-1000ms, the *unexpected plausible* and *implausible* words produced larger responses than the *expected* words, and the topographic sensor maps revealed distinct patterns for each effect, see Figure 4B.

**Figure 4.**
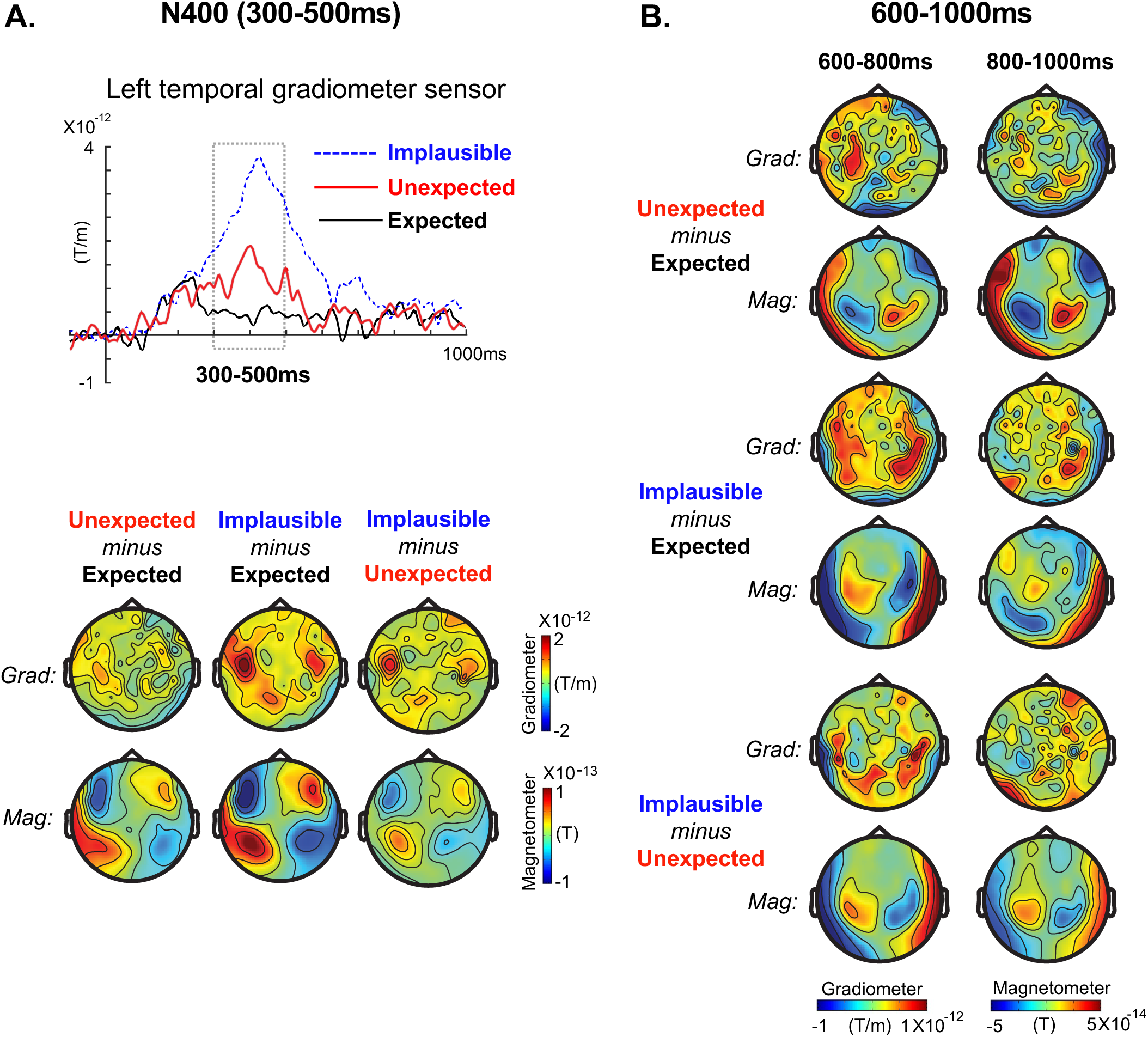
MEG sensor-level results. **A. 300-500ms.** *Top:* Grand-averaged event-related magnetic fields produced by critical words in each of the three conditions, shown at a left temporal gradiometer sensor (MEG0242+0243). The 300-500ms (N400) time-window is indicated using a dotted box. *Bottom:* MEG Gradiometer (Grad.) and Magnetometer (Mag.) sensor maps show the topographic distributions of the MEG N400 effects produced by contrasting the *Expected*, *Unexpected plausible* and *Implausible* critical words between 300-500ms. In all contrasts, the distribution of the MEG N400 effect was maximal over temporal sites, particularly on the left. **B. 600-1000ms.** MEG Gradiometer (Grad.) and Magnetometer (Mag.) sensor maps show the topographic distributions of the MEG effects produced by contrasting the *Expected*, *Unexpected plausible* and *Implausible* critical words in the first half (600-800ms) and the second half (800-1000ms) of the late time-window of interest. In order to better illustrate the scalp distribution of these late effects, these sensor maps are shown at a different scale from that used for the 300-500ms sensor maps. The contrasts between the *Unexpected plausible* and *Expected* critical words and the contrast between the *Implausible* and *Expected* critical words reveal somewhat distinct spatial distributions of sensor-level activity.

#### Source-localized MEG: Evoked effects

##### 300-500ms

As shown in Figure 5, between 300-500ms, *expected*, *unexpected plausible* and *implausible* words produced graded increases in activity within multiple regions of left lateral, ventral and medial temporal cortices. In medial temporal cortices, the effects were also driven by a dipole going in the opposite direction to the *expected* words. In addition, the *implausible* words produced a larger response than both the *expected* and *unexpected plausible* words within left-IFC, as well as a larger response than the *expected* words within anterior cingulate cortex.

**Figure 5.**
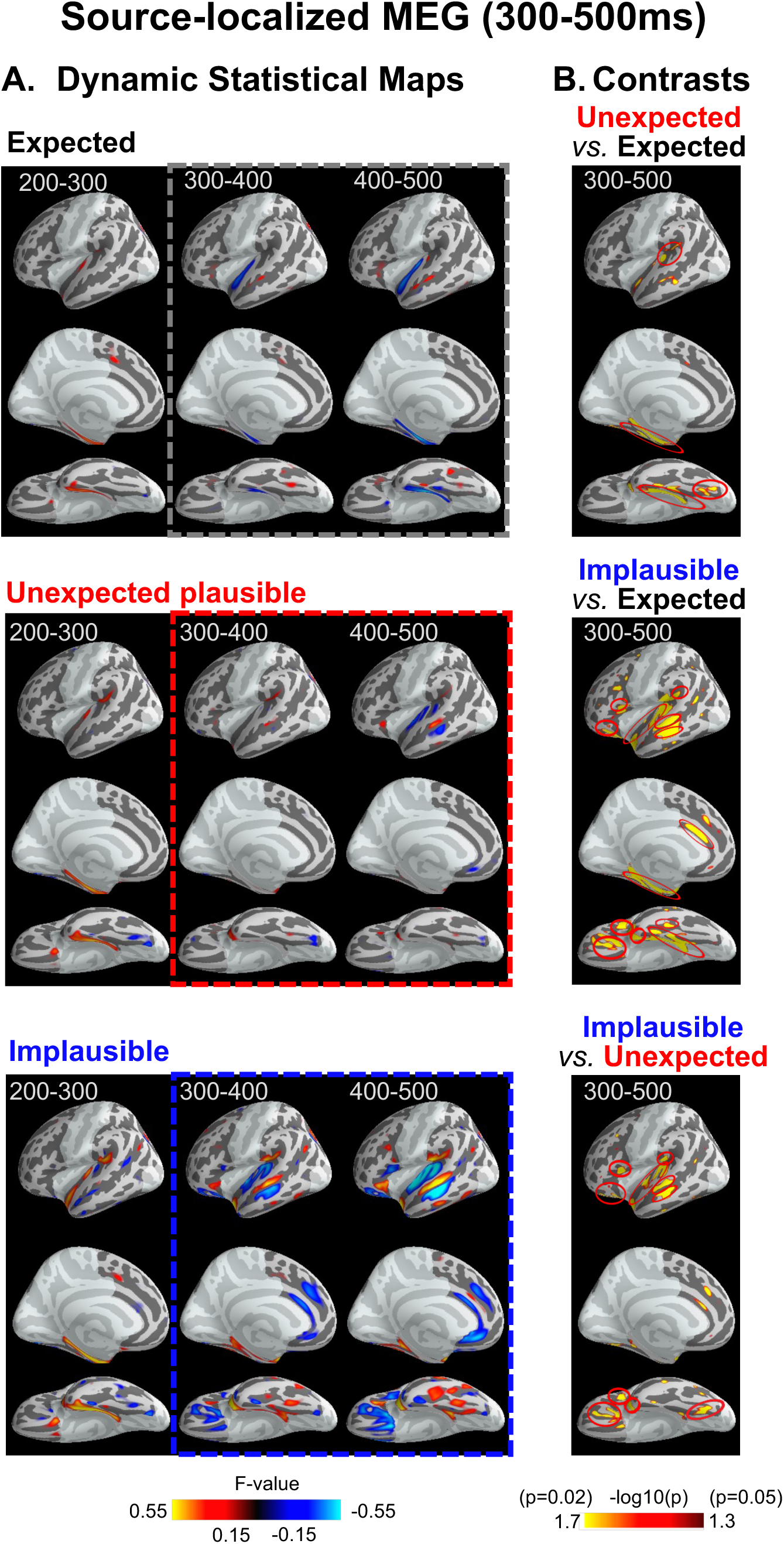
MEG source-level activity produced by the expected, unexpected plausible and implausible critical words in the 300-500ms (N400) time-window. **A.** Signed dynamic Statistical Parametric Maps (dSPMs) produced by *Expected* (*top*), *Unexpected plausible* (*middle*), and *Implausible* (*bottom*) critical words are shown at 100ms intervals from 200 until 500ms. All dSPMs are thresholded at 0.15, with red indicating outgoing currents (positive dSPM values) and blue indicating ingoing currents (negative dSPM values). Ingoing and outgoing currents that are directly adjacent to one another are interpreted as reflecting a single underlying dipole (neuroanatomical source). This is because if an underlying dipole/source is situated on one side of a sulcus, the signal can bleed into the other side, leading to the appearance of an adjacent dipole in the opposite direction^67^. The full dynamics of these source activations (at all sampling points) are shown as videos in Supplementary Materials. **B.** Statistical maps contrasting *Unexpected plausible* and *Expected* words (*top*), *Implausible* and *Expected* words (*middle*), and *Implausible* and *Unexpected plausible* words (*bottom*) within the 300-500ms (N400) time-window. Within left temporal cortex, these effects reached cluster-level significance, indicated using red circles, within (i) superior temporal gyrus, extending anteriorly towards the temporal pole and extending posteriorly into the supramarginal gyrus, (ii) the mid-portion of the superior temporal sulcus/middle temporal cortex (for contrasts involving the *Implausible* words), (iii) left ventral temporal cortex (mid- and posterior fusiform gyrus), and (iv) left medial temporal cortex (parahippocampal and entorhinal). The *Implausible* words additionally produced cluster-level effects in the left inferior frontal cortex (relative to both other conditions), and in the anterior cingulate cortex (relative to the *Expected* words). Both dSPMs and contrast maps are displayed on the FreeSurfer average surface, “fsaverage”^64^. Neuroanatomical regions were defined using the Desikan-Killiany Atlas^65^, see Supplementary Materials Figure 1 and Supplementary Table 1. See also Supplementary Materials for analyses over the right hemisphere (Supplementary Figures 2, 3 and 4).

##### 600-1000ms

As shown in Figure 6, by 500ms, the activity produced by the *unexpected plausible* words within left temporal cortex had diminished. Between 600-1000ms, however, they produced a response within left-IFC, and re-activated the left middle temporal cortex, with a dipole going in the opposite direction to that produced in the N400 time-window.

The evoked activity produced by the *implausible* words was quite different. By 500ms, the response they produced within the left-IFC had diminished, but the activity produced within left temporal and posterior fusiform cortices continued into the 600-1000ms time-window. In the latter half of this time-window, the *implausible* words also re-activated the left-IFC, with a dipole going in the opposite direction to that produced between 300-500ms. Finally, throughout the 600-100ms window, the *implausible* words also produced a large dipole within left medial temporal cortex, again with the opposite polarity to that produced in the N400 window.

Pair-wise statistical comparisons showed that, within left-IFC, there was a significant difference in comparing both the *unexpected plausible* and the *implausible* words with the *expected* words (600-800ms), with both effects being driven by dipoles going in opposite directions in the two conditions. A direct comparison between the *unexpected plausible* and *implausible* conditions, however, revealed no differences within left-IFC. Within left lateral temporal cortex, there were significant differences in comparing the *unexpected plausible* words with both the *expected* (800-1000ms) and *implausible* words (600-800ms). Both these effects were driven by dipoles going in the opposite direction in each of the two conditions. In posterior fusiform cortex there was a significant difference in comparing the *implausible* words with both the *expected* (600-800ms) and *unexpected plausible* words *(600-800ms; 800-1000ms).* Similarly, within medial temporal cortex, the response produced by the *implausible* words differed from that produced by both other conditions (600-800ms; 800-1000ms).

**Figure 6.**
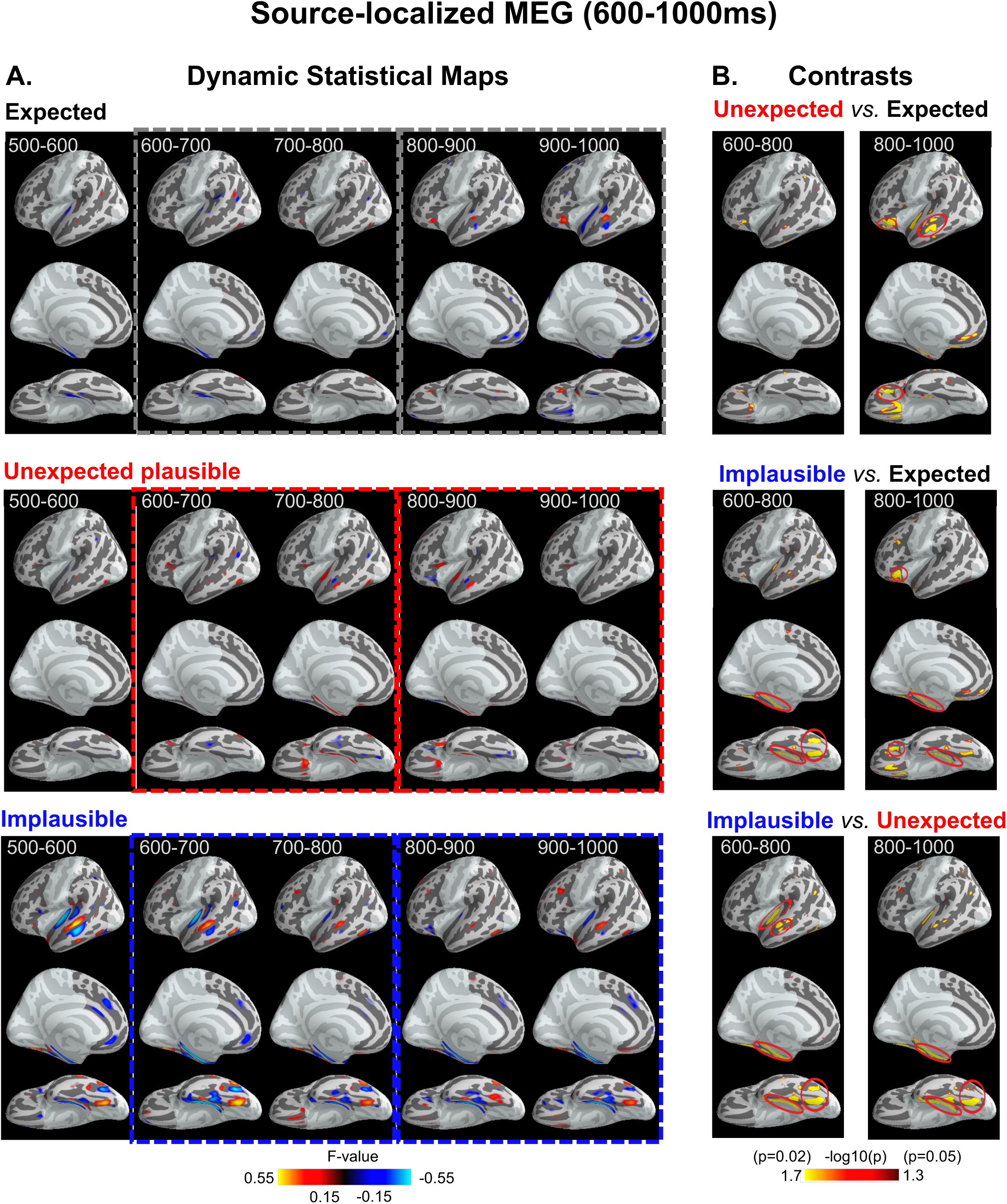
MEG source-level activity produced by the expected, unexpected plausible and implausible critical words in the late 600-1000ms time-window. **A.** Signed dynamic Statistical Parametric Maps (dSPMs) produced by *Expected* (*top*), *Unexpected plausible* (*middle*), and *Implausible* (*bottom*) critical words are shown at 100ms intervals from 500 until 1000ms. All dSPMs are thresholded at 0.15, with red indicating outgoing currents (positive dSPM values) and blue indicating ingoing currents (negative dSPM values). Ingoing and outgoing currents that are directly adjacent to one another are interpreted as reflecting a single underlying dipole (neuroanatomical source). This is because, if an underlying dipole/source is situated on one side of a sulcus, the signal can bleed into the other side, leading to the appearance of an adjacent dipole in the opposite direction^67^. The full dynamics of these source activations (at all sampling points) are shown as videos in Supplementary Materials. **B.** Statistical maps contrasting the *Unexpected plausible* and *Expected* words (*top*), the *Implausible* and *Expected* words (middle), and the *Implausible* and *Unexpected plausible* words (*bottom*) are shown between 600-800ms (*left*) and between 800-1000ms (*right*). Red circles indicate activity that reached cluster-level significance for each contrast. The *Unexpected plausible* versus *Expected* contrast revealed significant clusters within left middle temporal and inferior frontal cortices (driven by dipoles going in opposite directions in the two conditions). The *Implausible* versus *Expected* contrast revealed significant clusters within left posterior (occipitotemporal) fusiform cortex (driven by a dipole to the *Implausible* words), left inferior frontal cortex (driven by dipoles going in opposite directions in the two conditions), and within the medial temporal cortex (driven by a dipole to the *Implausible* words). Both dSPMs and contrast maps are displayed on the FreeSurfer average surface, “fsaverage”^64^. Neuroanatomical regions were defined using the Desikan-Killiany Atlas^65^, see Supplementary Materials Figure 1 and Supplementary Table 1. See also Supplementary Materials for analyses over the right hemisphere (Supplementary Figures 2, 3 and 4).

#### Source-localized MEG: Functional connectivity analysis

As shown in Table 2, between 300-500ms, activity to the *unexpected plausible* (versus *expected*) words, and to the *implausible* (versus *expected*) words within left lateral temporal cortex was functionally connected to left-IFC (above-chance cross-region classification). Between 600-1000ms, activity to the *unexpected plausible* (versus *expected*) words within left-IFC was functionally connected to left lateral temporal cortex. However, activity to the *implausible* (versus *expected*) words within left-IFC was not functionally connected either to left lateral temporal or posterior fusiform cortices.

**Table 2.** MEG Functional connectivity analysis: Cross-regional classifications.

| **Paired Regions of Interest** | **Classification Conditions** | **Accuracy**<br>**Mean (SD)** | ***t*-value**<br>***df* (31)** | ***p*-**<br>**value** | **Effect Size**<br>**(*d*)** |
| --- | --- | --- | --- | --- | --- |
| 300-500ms:<br>Left lateral temporal/<br>Left inferior frontal | Unexpected vs. Expected | 51.8% (5.2%) | 2.14 | 0.041 | 0.77 |
|  | Implausible vs. Expected | 52.9% (4.9%) | 3.27 | 0.003 | 1.17 |
| 600-1000ms:<br>Left inferior frontal/<br>Left lateral temporal | Unexpected vs. Expected | 52.0% (4.9%) | 2.12 | 0.042 | 0.76 |
|  | Implausible vs. Expected | 50.5% (6.0%) | 0.44 | 0.661 | 0.16 |
| 600-1000ms:<br>Left inferior frontal/<br>posterior fusiform | Unexpected vs. Expected | 49.4% (4.6%) | -0.77 | 0.447 | -0.27 |
|  | Implausible vs. Expected | 49.4% (5.3%) | -0.66 | 0.512 | -0.23 |

## Discussion

We used MEG and ERPs to track the spatiotemporal dynamics of activity evoked by *expected*, *unexpected plausible*, and *implausible* words during language comprehension. Our scalp-recorded ERP findings replicate previous studies by showing that, between 300-500ms, the three conditions produced a progressively larger N400^18, 19^, and between 600-1000ms, the *unexpected plausible* and *implausible* continuations produced spatially-distinct late positivities^18, 31, 32^. By simultaneously collecting MEG data and source-localizing this activity, we were able to show, for the first time, not only *when*, but also *where* these evoked effects were produced as information flowed up and down the left-lateralized fronto-temporal hierarchy during incremental comprehension. While several MEG^24, 25^ and intracranial studies^23^ have reported effects of linguistic context within temporal and/or prefrontal cortices within the N400 time-window, no previous study has distinguished between the effects of lexical predictability in plausible sentences, and the effects of implausibility. In addition, no previous study has systematically source-localized activity past the N400 time-window. By determining precisely when and how lower and higher levels of the fronto-temporal hierarchy are modulated following the onset of incoming words, our findings shed new light on the cognitive architecture and neurobiology of online language comprehension.

### 300-500ms

Consistent with previous ERP studies, the scalp-recorded N400 was larger to *unexpected plausible* than to *expected* words^13, 17^, and larger to *implausible* than to *unexpected plausible* words^18, 19^. The effect of implausibility was smaller in ERP than MEG (with implications for understanding the role of component overlap in interpreting ERP data, see Supplementary Materials). Within this 300-500ms time-window, our source-localized MEG data revealed graded effects within left temporal cortices (*Expected* < *Unexpected* < *Implausible*), but non-graded effects within left-IFC (*Implausible* > *Unexpected* = *Expected*).

#### Graded effects within left temporal cortex

Within left temporal cortex, the progressive increase in activity across the three conditions was observed in regions that are known to support lexico-semantic processing. These included left mid-fusiform^39, 40^ and superior temporal cortices^41, 42^, which function to map orthographic and phonological representations on to lexical representations; left middle temporal cortex, which is implicated in mapping between lexical and semantic representations^11^, and left anterior and medial temporal cortices, which may play a role in retrieving and “binding” distributed semantic features into distinct concepts^43^. We therefore take the graded effects of context within these regions as evidence of top-down lexico-semantic facilitation.

This account is consistent with predictive frameworks of language comprehension. According to such frameworks, predictions based on stored real-world and linguistic knowledge, are propagated down from higher to lower levels levels of representation, thereby facilitating the retrieval/access of the lexico-semantic features of anticipated incoming words^2, 11, 15, 17, 18, 26, 27, 29^. Therefore, when new bottom-up input becomes available to left temporal cortex, it is easiest to retrieve lexico-semantic representations of *expected* words, more difficult to retrieve those of *unexpected plausible* words^17^, and most difficult to retrieve those of *implausible* words whose semantic features were not predicted at all^18^.

Interestingly, the effects within the medial temporal cortex were not only driven by dipoles to the *unexpected plausible* and *implausible* continuations, but also by a dipole of the opposite polarity to the *expected* words. The precise significance of the dipole produced by the expected words is unclear. However, previous studies have shown that, in constraining contexts, medial temporal regions can be pre-activated prior to the onset of new input^44, 45^, and so we speculate that it indexed the detection of a match (so-called “resonance”^46^) between this pre-activated activity and the bottom-up input. This observation is consistent with previous intracranial studies showing that different electrodes within medial temporal regions produce local field potentials of opposite polarities to incongruous and expected words in the N400 time-window^23^, and it has important methodological implications for MEG analysis (see Supplementary Materials).

#### Non-graded effects within left inferior frontal cortex

In contrast to the graded increases in activity across the three conditions within left temporal cortex, the effects produced within left-IFC were not graded: in the N400 time-window, only the *implausible* words produced an enhanced evoked response within this region. This finding is more difficult to explain within existing frameworks of language comprehension. A fully predictive framework, which assumes all contextual effects “bottom-out” at the lexical level^26, 27^, would predicted no additional modulation within left-IFC. A single-state model, which posits continuous feedforward/feedback activity between left temporal and frontal cortices^29^, would predict graded effects within left-IFC (just as in left temporal cortex). A prediction-integration framework that attributes the effects of implausibility within the N400 time-window to higher-level “integration”^19, 20^ can explain the non-graded effects in left-IFC. However, this framework would incorrectly predict non-graded effects within left temporal cortex (lexico-semantic facilitation only in plausible sentences).

#### A predictive coding account of evoked activity within the N400 time-window

We suggest that the full pattern of evoked activity produced between 300-500ms can be explained by the principles of hierarchical predictive coding (see Figure 1). Like other top-down predictive frameworks, predictive coding posits that event representations that are being incrementally inferred within the left-IFC generates top-down lexico-semantic predictions that pre-activate upcoming lexico-semantic features. In contrast to these previous frameworks, however, instead of attributing the evoked response to the *change in state* that is induced by the arrival of new inputs at a given level of the cortical hierarchy^14, 28, 30^, predictive coding attributes the phase-locked evoked response to *prediction error*^5^: new bottom-up information that cannot be explained/suppressed by predictions from higher cortical areas^3–6^. Within this framework, the evoked response produced within left temporal cortex between 300-500ms is attributed to *lexico-semantic prediction error*––bottom-up lexico-semantic information that cannot be explained by top-down predictions. Therefore, like other predictive processing frameworks, predictive coding correctly predicts a progressively larger evoked response across the three conditions within left temporal regions: the less a newly-inferred lexico-semantic representation is suppressed by top-down predictions, the larger the lexico-semantic prediction error it produces.

Like single-state frameworks^29^, predictive coding also posits a continuous feedforward flow of information from left temporal to left-IFC throughout the N400 time-window. In predictive coding, this involves passing up the lexico-semantic prediction error to update higher-level event representations being inferred within left-IFC. And indeed, our functional connectivity analysis showed that, between 300-500ms, activity within left lateral temporal cortex produced by both the *unexpected plausible* and *implausible* (versus *expected*) words was functionally connected to left-IFC.

In contrast to single-state models, however, predictive coding correctly predicts that the evoked response produced by left-IFC is *non-graded* across the three conditions (*Implausible* > *Unexpected Plausible* = *Expected*). As Rao and Ballard explain in their original description of predictive coding in the visual system^4^, at higher levels of cortex, a robust prediction error is only produced by stimuli “whose statistics differ in certain drastic ways from natural statistics”. This is because higher-level regions themselves continually receive predictions, based on stored real-world knowledge, that suppress the higher-level prediction error produced by states that are congruous with these predictions. Therefore, when an update of the event model within left-IFC yields a *plausible* interpretation, event-level prediction error within this region is suppressed. However, when an update yields a highly *implausible* interpretation, higher-level prediction error is no longer suppressed, resulting in an enhanced evoked response within left-IFC.

Predictive coding also posits that the difficulty in converging on an implausible interpretation between 300-500ms within left-IFC results in less accurate top-down lexico-semantic predictions that therefore fail to “switch off” lower-level lexico-semantic prediction error within left temporal cortex. This failure to suppress lexico-semantic prediction error may have also contributed to the enhanced response to the *implausible* words within left temporal cortex within between 300-500ms, as well to the prolongation of this effect into the later 600-1000ms window, as discussed further below.

### 600-1000ms

Replicating previous ERP findings, the *plausible* and *implausible* words produced two distinct late positivities between 600-1000ms: a *late frontal positivity* to *unexpected plausible* words^18, 31–34^, and a *late posterior positivity/P600* to highly *implausible* continuations^18, 31–33, 35, 36^. Once again, the MEG source-localized results constrain our understanding of the functional significance of this late-stage evoked activity.

#### Late effects within left inferior frontal and middle temporal cortices to unexpected plausible words

It has been proposed that the *late frontal positivity* evoked by *unexpected plausible* words reflects high-level discourse shifts^20^, together with feedback activity to lower-level lexico-semantic representations^18, 33^. Our MEG findings support this account. Between 600-1000ms, the *unexpected plausible* words produced activity within left-IFC as well as a re-activation of the left middle temporal cortex. We suggest that these late effects reflected feedback activity that drove top-down shifts of the event model and lexico-semantic state.

Consistent with this top-down feedback account, our functional connectivity analysis revealed evidence of connectivity between left inferior frontal and temporal cortices when processing the *unexpected plausible* (versus *expected*) words in this late 600-1000ms time-window. In addition, the late re-activation of the left middle temporal cortex was driven by a dipole with the opposite polarity to that produced in the earlier N400 time-window. While the precise significance of this dipole reversal is unclear, it suggests that this late effect was functionally distinct from the earlier feedforward activity (lexico-semantic prediction error) produced between 300-500ms.

This account of the late fronto-temporal effect produced by the *unexpected plausible* inputs also follows naturally from the computational principles of a dynamic generative framework that assumes that our environments are “non-stationary”, changing rapidly from one situation to another^47^. According to this framework, unexpected inputs that violate prior high-confidence beliefs lead the brain to infer that the environment has systematically changed, initiating a “model shift” to a new “latent cause” that can better explain future inputs. For example, consider the scenario in which the event, <lifeguards warned trainees> was inferred in the N400 time-window (Figure 1). Even though this event is plausible, it is nonetheless out of keeping with the beach-related schema that had previously been inferred with high confidence, based on the prior discourse context. We suggest that this led comprehenders to infer an “event boundary”^48^, which initiated the retrieval of new schema-relevant information from long-term memory^37, 49^.

According to hierarchical predictive coding, this newly-retrieved schema then generated new top-down predictions that were passed down the cortical hierarchy via feedback connections (see Figure 2, left). Because these predictions encoded information that was not already represented within previously-inferred states, they produced “top-down error”^4^, both at the event and lexico-semantic levels. We suggest that the late evoked effect within left-IFC reflected top-down error at the event level, which drove the top-down shift in the event model. We also suggest that the late re-activation of left middle temporal cortex reflected top-down error at the lexico-semantic level, and that this drove a top-down shift to new schema-relevant lexico-semantic representations. This type of later-stage top-down re-activation has been described in the visual system when higher-level regions, encoding global gestalt representations, provide feedback enhancement of activity at lower-level regions that encode local image statistics^50^. During language comprehension, such dynamic feedback activity may play a critical role in ensuring that comprehenders continue to predict new inputs that are consistent with a newly-updated event model (e.g. new inputs related to what trainees might be doing on a beach).

#### Late effects within left posterior fusiform, inferior frontal and medial temporal cortices to highly implausible words

In the present study, the *implausible* words were not simply implausible—they were also anomalous (e.g., “*cautioned the *drawers”*); that is, they conflicted with the structure and state of the generative model as a whole. The detection of conflict may explain why the anterior cingulate cortex was activated in the earlier N400 time-window^51^.

It has been proposed that the *late posterior positivity/P600* produced by highly *implausible*/anomalous continuations between 600-1000ms reflects a conflict-driven reanalysis of the input at lower levels of linguistic representation^18, 33, 35^. Consistent with this account, the *implausible* words produced a robust late evoked effect within the posterior (occipitotemporal) fusiform cortex––the so-called “visual word-form area” that supports sublexical orthographic processing^9, 10^.

Predictive coding provides an account of this orthographic reanalysis (see Figure 2, right). Within this framework, evoked activity within left posterior fusiform cortex is attributed to the generation of orthographic prediction error at a still lower level of linguistic representation^9^. We suggest that this late orthographic prediction error was produced because higher levels of cortex failed to converge on a single, stable interpretation between 600-1000ms, resulting in a failure to generate accurate top-down predictions that suppressed the low-level error. Specifically, in contrast to the *unexpected plausible* inputs, it was not possible to retrieve new stored schemas from long-term memory to explain the highly implausible event model inferred between 300-500ms (<lifeguards cautioned drawers>). Therefore, between 600-1000ms, predictions based on the prior context and real-world knowledge continued to be propagated down the cortical hierarchy. Within left middle temporal cortex, these top-down lexico-semantic predictions (e.g., “swimmers” <animate>) would have been incompatible with the lexico-semantic information that was inferred from the bottom-up input (“drawers” <inanimate>), resulting in a destabilization of the lexico-semantic state. We suggest that, as a result of this destabilization, the left middle temporal cortex produced noisy, inaccurate orthographic predictions that were passed down via feedback connections to the posterior fusiform cortex^52^, leading to a failure to suppress orthographic prediction error, and to an enhanced evoked response within this region (orthographic reanalysis).

We further suggest that the re-activation of left IFC (with a dipole of the opposite polarity to that produced in the N400 time-window) was driven by top-down event error, produced when the top-down event predictions (<lifeguards cautioned swimmers>) failed to explain the implausible event that had been inferred within the N400 time-window (<lifeguards cautioned drawers>). The presence of conflicting information across multiple levels of representation explains why, in this later 600-1000ms time-window, there was no evidence of functional connectivity between the left-IFC and either left lateral temporal or posterior fusiform cortices to the *implausible* (versus *expected*) words.

Finally, the *implausible* continuations also produced a dipole within the medial temporal cortex throughout the 600-1000ms time-window, again with the opposite polarity to that produced in the N400 time-window. We speculate that this medial temporal activity supported new learning/adaptation, which was triggered by the detection of strong prediction violations^53, 54^, and the failure of the current generative model to explain the input^37^––that is, to minimize error across the cortical hierarchy. This interpretation would be consistent with growing evidence for a computational role of prediction error in bridging language processing and learning^55^, as well as with known links between the *late posterior positivity/P600* ERP component and adaptation to new environments^56^.

## Conclusions

We have argued that the full time-course of evoked activity produced by *expected*, *unexpected plausible* and *implausible* words across left temporal and inferior frontal cortices can be understood within a hierarchical predictive coding framework of language comprehension. This interpretation leaves open many questions that will be important to address in future studies. Consistent with a predictive coding architecture, we have assumed that the phase-locked evoked response specifically indexes prediction error^5^, which is functionally distinct from the process of updating state representations^4, 5^. It will therefore be important to determine whether multivariate methods can detect implicit state updates that can be dissociated from the phase-locked evoked response. It will also be important for future studies to parametrically manipulate lexical predictability and implausibility to determine precisely how these factors modulate evoked activity within left temporal and inferior frontal cortices at both early and later stages of processing.

Of course, no single study can provide definitive evidence for any single model of language comprehension. As we have noted, several of our individual findings, such as the graded increases in activity across the three conditions within left temporal cortex in the N400 time-window, or the reanalysis within the posterior fusiform cortex to the highly implausible inputs in the later 600-1000ms time-window, are also consistent with other psycholinguistics or neurobiological frameworks that have attempted to explain the corresponding ERP effects at the scalp surface. Here, we have interpreted the full pattern of findings within a single computational framework that has been proposed as unifying theory of brain function, across multiple domains of perception and cognition^6^, including lower-level aspects of language processing^7–10^. Our findings suggest that the computational principles of predictive coding may also explain the flow of bottom-up and top-down information across the left-lateralized fronto-temporal network that supports higher-level language comprehension.

## Methods

### Materials

Participants read three types of three-sentence discourse scenarios, each with a constraining context, and a critical noun in the third sentence: (1) *Expected*, in which the critical word was predictable; (2) *Unexpected plausible*, in which the critical word was plausible but unpredictable because it was out of keeping with the schema set up by the prior context, and (3) *Implausible*, in which the critical word violated the animacy constraints of the preceding verb (which constrained either for an animate or an inanimate noun).

The stimuli were based on three of the conditions used in a previous ERP study^18^. A full description is provided there as well as in Supplementary Materials. Briefly, the discourse contexts of each scenario were constraining (average cloze probability of the most probable word: 68%), as quantified in a cloze norming study that was carried out in participants recruited through Amazon Mechanical Turk. These contextual constraints came from the entirety of the discourse context—the first two sentences plus the first few words of the third sentence before the critical word. In all scenarios, these first few words of the third sentence included an adjunct phrase of 1-4 words, followed by a pronominal subject that referred back to the first two sentences, a verb and a determiner. The verb was always relatively non-constraining in minimal contexts (cloze probability of the most probable word was below 24%, as quantified in another cloze norming study in which participants, recruited through Amazon Mechanical Turk, were presented with just a proper name, the verb, and a determiner).

To create the *expected* scenarios, each context was paired with the noun with the highest cloze probability for that context. To create the *unexpected plausible* scenarios, each context was paired with a noun of zero (or very low) cloze probability, but that was still plausible in relation to this context. To create the *implausible* scenarios, each context was paired with a noun that violated the animacy constraints of the preceding verb, see Table 3. In all scenarios, the critical noun was followed by three additional words to complete the sentence.

**Table 3.** Example of each experimental condition, together with stimuli characteristics.

| **Prior discourse-constraining context:** *The lifeguards received a report of sharks right near the beach. Their immediate concern was to prevent any incidents in the sea. Hence, they cautioned the...* |  |  |  |  |  |  |  |  |
| --- | --- | --- | --- | --- | --- | --- | --- | --- |
| **Scenario Type** | **Critical word** | **\*Constraint** | **\*\*Cloze** | **+SSV** | **Length** | **++Freq.** | **^OLD** | **^^Conc.** |
| Expected | <u>swimmers...</u> | 69%<br>(14%) | 69%<br>(14%) | 0.18<br>(.18) | 5.69<br>(1.60) | 1.53<br>(0.66) | 1.93<br>(0.56) | 4.30<br>(0.69) |
| Unexpected<br>Plausible | <u>trainees...</u> | 69%<br>(14%) | 0.1%<br>(0.5%) | 0.01<br>(.06) | 7.46<br>(2.22) | 0.61<br>(0.88) | 2.61<br>(0.86) | 4.15<br>(0.69) |
| Implausible | <u>drawers...</u> | 66%<br>(16%) | 0%<br>(0%) | 0.01<br>(.05) | 7.11<br>(2.04) | 0.81<br>(0.85) | 2.47<br>(0.81) | 4.21<br>(0.66) |

The critical words (underlined here although not in the experiment itself) were followed by three additional words, indicated here with three dots.
Means are shown with the standard deviations in parentheses.
\*The lexical constraint of each discourse context was calculated by identifying the most common completion across participants who saw that context in a cloze norming study (see Supplementary Materials), and tallying the proportion of participants who provided this completion.
\*\*Cloze probabilities of critical words were calculated based on the percentage of respondents in the cloze norming study who provided the critical noun.
+SSV: Semantic Similarity Values, quantifying the semantic relatedness between the critical words and the “bag of words” within the prior contexts, based on Latent Semantic Analysis.
Length: Number of letters.
++Freq.: Log Frequency values of critical words, retrieved from the English Lexicon Project.
^OLD: Orthographic Levenshtein Distance values of critical words, retrieved from the English Lexicon Project.
^^Conc.: Concreteness ratings of critical words<sup>70</sup>.

Each list contained 25 *expected*, 25 *unexpected plausible,* and 50 *implausible* scenarios. This ensured that each participant saw an equal proportion of plausible and implausible scenarios. Conditions were counterbalanced across lists such that, across all participants, (a) the same 25 discourse contexts appeared in all three conditions, but no individual saw the same context more than once, and (b) the same 25 critical words appeared in the 25 *unexpected plausible* scenarios and 25 of the *implausible* scenarios, but no individual saw this word more than once. We made the *a priori* decision to include all 50 implausible scenarios in our main analyses in order to maximize power to source-localized effects (see below). As a byproduct of counterbalancing the same high constraint contexts across the three conditions, the critical words in the *expected* scenarios had fewer letters, smaller orthographic neighborhoods and were more frequent than in the two other conditions (all *ts* > 5, *ps* < 0.001, *ds* > 0.54). However, all these lexical features were matched between the *unexpected plausible* and the *implausible* conditions (all *ts* < 1.74, *ps* > 0.08, *ds* < 0.16), see Table 3. In addition, semantic relatedness between the critical words and their prior contexts (operationalized using Semantic Similarity Values, SSVs, extracted using Latent Semantic Analysis^57^, http://lsa.colorado.edu/, term-to-document with default settings) were matched between the *unexpected plausible* and *implausible* scenarios (*t_(248)_* = 0.09, *p* = 0.93, *d* = 0.01). As discussed in Supplementary Materials, each list additionally included 100 scenarios with less constraining contexts (again, 50% plausible and 50% implausible). The 50 *low constraint implausible* scenarios served as fillers, and the 50 *low constraint plausible* scenarios served as a fourth condition. Analyses involving this fourth condition will be reported in a separate manuscript.

### Overall procedure

Participants took part in two separate experimental sessions: one in which we simultaneously collected MEG/EEG data, and one in which we collected structural and functional MRI data. The present manuscript focuses on the EEG and MEG datasets. Below, we describe the acquisition and analysis of the MEG/EEG data, as well as the structural MRI data, which was used to constrain MEG source localization. A report of the fMRI dataset, together with detailed comparisons with the EEG/MEG dataset, will appear in a separate manuscript.

### Participants

Thirty-three participants took part, but one MEG/EEG dataset was excluded because of technical problems. Here we report the results of 32 MEG/EEG datasets (16 females, mean age: 23.4; range: 18-35). All participants were native speakers of English (with no other language exposure before the age of 5), were right-handed, and had normal or corrected-to-normal vision. All were screened to exclude past or present psychiatric and neurological disorders, and none were taking medication affecting the Central Nervous System. The study was approved by the Mass General Brigham Institutional Review Board (IRB), and written informed consent was obtained from all participants.

### Stimuli presentation and task

Stimuli were presented using PsychoPy 1.83 software and projected on to a screen in white Arial font (size: 0.1 of the screen height) on a black background. On each trial, the first two sentences appeared in full (each for 3900ms, 100ms interstimulus interval, ISI), followed by a fixation (a white “++++”), which was presented for 550ms, followed by a 100ms ISI. Then the third sentence was presented word by word (each word for 450ms, 100ms ISI).

Participants’ task was to judge whether or not the scenario “made sense” by pressing one of two buttons after seeing a ‘‘?’’, which appeared after each scenario (1400ms with a 100ms ISI). Response fingers were counterbalanced across participants. This task encouraged active coherence monitoring during online comprehension and was intended to ensure that comprehenders detected the anomalies, which is necessary to produce a neural response^58^. In addition, following approximately 24/200 trials (distributed semi-randomly), participants answered a “YES/NO” comprehension question that appeared on the screen for 1900ms (100ms ISI). This encouraged participants to comprehend the scenarios as a whole, rather than focusing on only the third sentence in which the anomalies appeared.

Following each trial, a blank screen was presented with a variable duration that ranged from 100-500ms. This was then followed by a green fixation (++++) for a duration of 900ms, followed by an ISI of 100ms. These green fixations were used to estimate the noise covariance for the MEG source localization (see below). To ensure precise time-locking of stimuli, we used frame-based timing, which synced stimulus presentation to the frame refresh rate of the monitor (for example, a 450ms word presentation would be displayed for exactly 27 frames on our 60Hz monitor).

Stimuli were presented in eight blocks, each with 25 scenarios. Blocks were presented in random order in each participant. Participants took part in a short practice session before the formal experiment to gain familiarity with the stimulus presentation and tasks.

### Data acquisition

#### MEG and EEG data acquisition

Participants sat inside a magnetically shielded room (IMEDCO AG, Switzerland). The MEG data were acquired with a Neuromag VectorView system (Elekta-Neuromag Oy, Finland) with 306 sensors—102 triplets, with each triplet comprising two orthogonal planar gradiometers and one magnetometer. The EEG data were acquired at the same time using a 70-channel MEG-compatible scalp electrode system (BrainProducts, München), and referenced to an electrode placed on the left mastoid. An electrode was also placed on the right mastoid and a ground electrode was placed on the left collarbone. EOG data were collected with bipolar recordings: vertical EOG electrodes were placed above and below the left eye, and horizontal EOG electrodes were placed on the outer canthus of each eye. ECG data were also collected with bipolar recordings: ECG electrodes were placed a few centimeters under the left and right collarbones. Impedances were kept at <20kΩ at all scalp sites, at <10kΩ at mastoid sites, and at <30kΩ at EOG and ECG sites. Both MEG and EEG data were acquired with an online band-pass filter of 0.03-300Hz and were continuously sampled at 1000Hz.

To record the head position relative to the MEG sensor array for later co-registration of the MEG and MRI coordinate frames, the locations of three fiduciary points (nasion and two auricular), four head position indicator coils, all EEG electrodes, and at least 100 additional points, were digitized using a 3Space Fastrak Polhemus digitizer, integrated with the Vectorview system.

#### Structural MRI data acquisition

In order to create participant-specific head model for MEG source localization, we acquired structural MRIs using a 3T Siemens Trio scanner with a 32-channel head coil for all participants. T1-weighted high-resolution structural images were obtained using the following parameters: 1mm isotropic multi-echo magnetization-prepared rapid gradient-echo, MP-RAGE; Time to Repetition (TR): 2.53s; flip angle: 7 degrees; 4 echoes with TE: 1.69ms, 3.55ms, 5.41ms, and 7.27ms.

### ERP Analysis

EEG data were analyzed using the Fieldtrip software package^59^ in the Matlab environment. EEG channels with excessive noise (7 out of the 70 channels, on average) were visually identified and marked as bad channels. We then applied a low band-pass filter (30Hz), down sampled the EEG data to 500Hz, and segmented the epochs from -2600ms to 1400ms, relative to the onset of the critical words. After that, we visualized the data in summary mode within the Fieldtrip toolbox to identify the trials that showed high variance across channels. These trials were then removed from subsequent analysis. We then carried out an Independent Component Analysis (ICA) to remove ICA components associated with eye-movement (one component on average was removed per participant). Finally, we visualized the artifact-corrected trials, and removed any additional trials with residual artifact. On average, 6.5% of trials were removed from each condition (equally distributed across the three conditions: *F*(2,62) = 1.40, *p* = 0.26, *η^2^* = 0.048), yielding, on average, 23 trials in the *expected* and *unexpected plausible* conditions, and 46 trials in the *implausible* condition. Finally, the data of bad channels were interpolated using spherical spline interpolation^60^. In each participant, at each site, we then calculated ERPs, time-locked to the onset of critical words, in each of the three conditions, applying a -100ms pre-stimulus baseline.

We next averaged these voltages across all time points and electrode sites within each of three spatiotemporal regions of interest (ROIs) that were selected, *a priori,* to capture the N400, the *late frontal positivity* and the *late posterior positivity/P600* ERP components^18^. The N400 was operationalized as the average voltage across ten electrode sites within a central region (Cz, C1, C2, C3, C4, CPz, CP1, CP2, CP3, CP4), averaged across all sampling points between 300-500ms; the *late frontal positivity* was operationalized as the average voltage across eight electrode sites within a prefrontal region (FPz, FP1, FP2, FP3, FP4, AFz, AF3, AF4), averaged across sampling points between 600-1000ms; the *late posterior positivity/P600* was operationalized as the average voltage across 11 electrode sites within a posterior region (Pz, P1, P2, P3, P4, POz, PO3, PO4, Oz, O1, O2), averaged across sampling points between 600-1000ms. We carried out planned statistical comparisons between each pair of conditions. We also report analyses across an earlier 200-300ms time-window in Supplementary Materials.

### MEG Analysis

#### MEG preprocessing, individual averaging and sensor-level visualization

MEG data were analyzed using version 2.7.4 of the Minimum Norms Estimate (MNE) software package in Python^61^. In each participant, in each run, MEG sensors with excessive noise were visually identified and removed from further analysis. This resulted in the removal of seven (on average) of the 306 MEG sensors. Signal-space projection (SSP) correction was used to correct for ECG artifact. Trials with eye-movement and blink artifacts were automatically removed^61^. Then, after applying a band-pass filter at 0.1Hz to 30Hz, we segmented epochs from -100 to 1000ms, relative to the onset of the critical words. We removed epochs with additional artifact, as assessed using a peak-to-peak detection algorithm (the pre-specified cutoff for the maximal amplitude range was 4×10^-10^ T/m for the gradiometer sensors and 4×10^-12^ T for the magnetometer sensors). On average, 6.7% trials in each condition were removed (equally distributed across the three conditions: *F*(2,62) = 0.87, *p* = 0.42, *η^2^* = 0.027), yielding, on average, 21 artifact-free trials in the *expected* and *unexpected plausible* conditions, and 42 artifact-free trials in the *implausible* condition.

In each participant, in each block, at each magnetometer sensor and at each of the two gradiometers at each site, we calculated event-related fields (ERFs), time-locked to the onset of critical words in each of the three conditions, applying a -100ms pre-stimulus baseline. We averaged the ERFs across blocks in sensor space, interpolating any bad sensors using spherical spline interpolation^60^. We created gradiometer and magnetometer sensor maps to visualize the topographic distribution of ERFs across the scalp. In creating the gradiometer maps, we used the root mean square of the ERFs produced by the two gradiometers at each site.

#### MEG source localization in individual participants

Each participant’s cortical surface was first reconstructed from their structural T1 MPRAGE image using the FreeSurfer software package developed at the Martinos Center, Charlestown, MA (http://surfer.nmr.mgh.harvard.edu). We used MNE-Python^61^ to estimate the sources of the ERFs evoked by critical words in each of the three conditions, on each participant’s reconstructed cortical surface using Minimum-Norm Estimation (MNE)^62^.

In order to calculate the inverse operator in each participant—the transformation that estimates the underlying neuroanatomical sources for a given spatial distribution of activity in sensor space—we first needed to construct a noise-covariance matrix of each participant’s MEG sensor-level data, as well as a forward model in each participant (the model that predicts the pattern of sensor activity that would be produced by all dipoles within the source space).

To construct the noise covariance matrix in each participant, we used 650ms of MEG sensor-level data recorded during the presentation of the green inter-trial fixations (we used an epoch from 100-750ms, which cut off MEG data measured at the onset and offset of these fixations in order to avoid onset and offset evoked responses). We concatenated these fixations across blocks. To construct the forward model in each participant, we needed to (a) define the source space—the location, number and spacing of dipoles, (b) create a Boundary Element Model (BEM), which describes the geometry of the head and the conductivities of the different tissues, and (c) specify the MEG-MRI coordinate transformation—the location of MEG sensors in relation to the head surface.

The source space was defined on the white matter surface of each participant’s reconstructed MRI and constituted 4098 vertices per hemisphere, with three orthogonally-orientated dipoles at each vertex (two tangential and one perpendicular to the cortical surface). We defined these vertices using a grid that decimated the surface into meshes, with a spacing of 4.9mm between adjacent locations (spacing: “oct6”). We created a single compartment BEM by first stripping the outer non-brain tissue (skull and scalp) from the pial surface using the watershed algorithm in FreeSurfer, and then applying a single conductivity parameter to all brain tissue bounded by the inner skull. We specified the location of the MEG sensors in relation to the head surface by manually aligning the fiducial points and 3D digitizer (Polhemus) data with the scalp surface triangulation created in FreeSurfer, using the mne_analyze tool^61^.

We then calculated the inverse operator in each participant, setting two additional constraints. First, we set a loose constraint on the relative weighting of tangential and perpendicular dipole orientations within the source space (loose = 0.2). Second, we set a constraint on the relative weighting of superficial and deep neuroanatomical sources (depth = 0.8) in order to increase the likelihood that the minimum norm estimates would detect deep sources.

We then applied each participant’s inverse operator to the ERFs of all magnetometer and gradiometer sensors calculated within each block. We estimated activity at the dipoles that were orientated perpendicular to the cortical surface at each vertex (pick_ori = “normal”). Each of these perpendicular dipoles had both a positive and a negative value, which indicated whether the currents were outgoing or ingoing respectively. We chose to retain the two polarities of each estimated dipole for further analyses for two reasons. First, this approach allowed us to include all trials in each of our three conditions, thereby maximizing power without inflating our estimate of noise in the conditions with more trials (if we had chosen to simply estimate the magnitude of each dipole by squaring the positive and negative values to yield positively-signed estimates, we would have artificially inflated the noise estimates in the *implausible* conditions, which had twice as many trials as the *expected* and the *unexpected plausible* conditions). Second, by retaining this polarity information, we were able to determine whether any statistical differences between conditions were driven by differences in the magnitude and/or differences in the polarity of the dipoles evoked in each condition (see Supplementary Materials for further discussion).

Then, for each condition in each block, we computed noise-normalized dynamic Statistical Parametric Maps (dSPMs)^63^ on each participant’s cortical surface at each time point, and averaged these values across blocks within each participant. Finally, the source estimates of each participant were morphed on the FreeSurfer average brain “fsaverage”^64^ for group averaging and statistical analysis.

#### Statistical analysis of MEG source-level evoked activity

To statistically analyze the source-localized evoked (phase-locked) MEG responses, we carried out planned comparisons between each pair of conditions over a large left-lateralized search region that included classic language-related areas as well as other regions of interest (left lateral temporal cortex, left ventral temporal cortex, left medial temporal cortex, left lateral parietal cortex, left lateral frontal cortex, and left medial frontal cortex). This search area was defined on the Desikan-Killiany Atlas^65^, and is illustrated in Supplementary Figure 1A. Within this search region, we examined activity within three 200ms time-windows of interest: 300-500ms, corresponding to the N400 time-window, and 600-800ms and 800-1000ms, corresponding to the first and second halves of the time-window associated with late positivity ERP effects. To account for multiple comparisons, we tested hypotheses using permutation-based cluster mass procedures^66^, which were modified as described next. Note that this non-parametric cluster-based approach is robust to any differences in signal-to-noise resulting from different numbers of trials in the *expected*, *unexpected plausible* and *implausible* conditions.

Within each 200ms time-window, we carried out pairwise t-tests between each pair of conditions on the signed estimated dSPM values at each vertex and at each time point. Instead of using the resulting signed t-values to compute our cluster-level statistic, we used unsigned -log-transformed p-values. This is because a single neuroanatomical source that is located on one side of a sulcus can appear on the cortical surface as adjacent dipoles of opposite polarity (outgoing and ingoing currents) because of signal bleeding to the other side of the sulcus^67^. This is apparent in the activation maps that show the signed dSPM values at each location in each condition (see Figures 5 and 6): positive dSPM values, corresponding to outgoing currents (shown in red), and negative dSPM values, corresponding to ingoing currents (shown in blue), often appear on either side of a sulcus, but these are likely to reflect a single underlying dipole/neuroanatomical source. The use of unsigned p-values therefore ensured that adjacent effects of opposite signs were treated as a single cluster/single underlying source in the statistical analyses. Within each time-window of interest, any data points that exceeded a pre-set uncorrected significance threshold of 1% (i.e., *p* ≤ 0.01) were -log10 transformed, and the rest were zeroed.

In order to account for multiple spatial comparisons, we subdivided the search area into 140 equal-sized patches^68^, shown in Supplementary Figure 1B. Within each patch, we took the average of the -log-transformed p-values across all time points within each time-window of interest (300-500ms, 600-800ms, 800-1000ms) as our cluster statistic. To compute a null distribution for the cluster-mass statistic, we carried out exactly the same procedure as that described above, but this time we randomly assigned dSPM values between the two conditions for a given contrast, and repeated this procedure 10,000 times. For each randomization, we took the largest value across all spatial patches as our cluster mass statistic. To test our hypotheses at each spatial patch in each time-window of interest, we compared the observed cluster-level statistic for that patch against this null distribution. If our observed cluster-level statistic fell within the highest 5.0% of the distribution, we considered it to be significant. Note that this approach allowed us to account for temporal and spatial discontinuities in effects (resulting from noise). However, it constrains any statistical inference to the spatial resolution of each patch and to the temporal resolution of our *a priori* time-windows.

In order to illustrate the results, we projected the averaged uncorrected -log10 transformed p-values (*p* < 0.05) at each vertex on to the “fsaverage” brain. We use circles to indicate any spatial patches in which we observed a significant cluster, grouping these areas by the anatomical regions shown in Supplementary Figure 1A and listed in Supplementary Table 1.

Finally, we carried out the following exploratory analyses, which are reported in Supplementary Materials: (a) an analysis of a subset of the MEG data to address potential concerns regarding differences between the *expected* and *implausible* conditions in the number of trials per condition, and in the lexical properties of the critical words, (b) an analysis to determine whether there were effects in an earlier 200-300ms time-window, and (c) an analysis of an analogous search region within the right hemisphere.

#### MEG connectivity analysis within spatial regions of interest

We carried out functional connectivity analyses using a cross-ROI multivariate decoding approach^69^ to determine whether, for particular contrasts of interest, a pair of regions that were co-activated within a particular time-window encoded shared information, i.e. whether they were functionally connected. For each of these analyses, we extracted single-trial MEG activity from individual vertices within two spatial ROIs that produced MEG evoked effects in source space (at p ≤ 0.05 uncorrected). For analyses in the 300-500ms time-window, we defined a left lateral temporal and a left inferior frontal ROI, based on the contrast between the *implausible* and *expected* words. For analyses in the 600-1000ms window, we defined a left lateral temporal and left inferior frontal ROI, based on the contrast between the *unexpected plausible* and *expected* words, and a left posterior fusiform and left inferior frontal ROI, based on the contrast between the *implausible* and *expected* words.

To extract the MEG data within each ROI, we applied each participant’s inverse operator to their single-trial sensor-level MEG data (at all magnetometer and gradiometer sensors), yielding a matrix with dimensions, *trial × vertex × time*. To ensure that activity within a pair of ROIs was in the same dimensional space, we rotated these matrices in the direction of maximum variance using Principal Component Analysis (PCA), keeping components that explained at least 99% of the variance across all time samples. We conducted connectivity analysis on these PCA-transformed data using the *sklearn* package in python. Using logistic regression, we first trained classifiers to distinguish activity between the two conditions that comprised a given contrast at each time point within one of the two ROIs, and then asked whether these trained classifiers were able to discriminate activity between the same two conditions at corresponding time points within its paired ROI. We averaged the cross-ROI decoding accuracy values across all time points within the time-window of interest and used a one-sample t-test to determine whether this average decoding accuracy was significantly different from 50% (chance level). A significant cross-ROI decoding performance indicates significant functional connectivity between the two ROIs.

## Acknowledgements

This work was funded by the National Institute of Child Health and Human Development (R01 HD082527 to G.R.K.). We thank Nao Matsuda for her technical MEG support, as well as Edward Wlotko, Nate Delaney-Busch, Eric Fields, Allison Fogel and Arim Choi Perrachione for their assistance with data collection. We also thank Seppo Ahlfors for his help with data analysis, and Jasmine Falk and Rebeca Becdach for their help in data visualization. This research was carried out at the Athinoula A. Martinos Center for Biomedical Imaging at the Massachusetts General Hospital, using resources provided by the Center for Functional Neuroimaging Technologies, P41EB015896, a P41 Biotechnology Resource Grant supported by the National Institute of Biomedical Imaging and Bioengineering (NIBIB), National Institutes of Health. This work also involved the use of instrumentation supported by the NIH Shared Instrumentation Grant Program, specifically, grant numbers S10RR014978 and S10RR021110. Finally, we thank Jeff Stibel for his support of Drs. Kuperberg and Wang.

## Author contributions statement

Experimental design: LW (Lin Wang), LS, GK; Data acquisition: LW (Lin Wang), LS, EA, LW (Lena Warnke), MK; Data analysis: LW (Lin Wang), LS, EA, SK, MH, GK; Data interpretation: LW (Lin Wang), LS, TB, GK; Manuscript preparation: LW (Lin Wang), LS, TB, GK.

