## Supplementary Materials for "Predictive coding across the left fronto-temporal hierarchy during language comprehension"


**Supplementary Methods**

1. Supplementary details about stimuli

*(a) Development of stimuli and cloze norming studies*

*(b) List composition and counterbalancing*

2. Spatial search region used for MEG source localization

**Supplementary Results**

1. Supplementary behavioral results

2. Exploratory analysis of ERP and source-localized MEG data between 200-300ms

3. MEG source-localization videos

4. Exploratory analysis of a subset of the MEG data

5. Exploratory analysis of MEG source-level activity over the right hemisphere

**Supplementary Discussion**

1. The relationship between the MEG and ERP findings

2. The retention of dipole polarity when carrying out MEG source localization

*(a) Different conditions can produce dipoles of opposite polarities within the time-windows and anatomical regions*

*(b) The same condition can produce dipoles within the same region that reverse in polarity across time-windows*

*(c) Methodological and theoretical implications*

### Supplementary Methods

1. Supplementary details about stimuli

*(a) Development of stimuli and cloze norming studies*

To develop the discourse scenarios, we carried out two cloze norming studies. For both norming studies, participants were recruited through Amazon Mechanical Turk. They were asked to complete each context with the first word that came to mind (Taylor, 1953), and in an extension of the standard cloze procedure, to then provide two additional words that could complete the sentence (Federmeier, Wlotko, De Ochoa-Dewald, & Kutas, 2007; Schwanenflugel & Lacount, 1988). Individuals were excluded if (a) their first language was anything other than English, (b) they self-reported any psychiatric or neurological disorders, or (c) they failed to follow instructions (we included "catch" questions that served as attention checks).

The first cloze norming study aimed to characterize a subset of verbs, which we used to construct the final sentences of each three-sentence scenario. Specifically, this norming study served to establish the lexical constraints and animacy constraints of these verbs in minimal contexts. We began with a large set of 617 transitively-biased verbs that were taken from a number of different sources, including a set of linguistically-characterized verbs (Levin, 1993) and materials from previous studies carried out in our lab (Paczynski & Kuperberg, 2011, 2012). We excluded verbs with a log Hyperspace Analogue to Language (HAL) frequency (Lund & Burgess, 1996) of two standard deviations below the mean (based on English Lexicon Project database, (Balota et al., 2007). For each verb, we constructed a minimum context, consisting of a proper name, the verb, and a determiner (e.g., *"Harry explored the…"*). For cloze norming, we divided these sentence stems into six lists in order to reduce time demands on any individual. After exclusions, between 89 and 106 participants provided completions for each item.

Based on the animacy of the noun completions, we categorized the verbs as either animate constraining or inanimate constraining, and tallied the number of participants who produced the best completions in order to calculate the lexical constraints of each verb in minimal contexts. We then selected a subset of these verbs (50% animate constraining; 50% inanimate constraining), with lexical constraints less than 24%.

For each verb, we then wrote a corresponding discourse context. Each discourse context consisted of two introductory sentences, and a third sentence that included an adjunct phrase (1-4 words), a pronominal subject that referred back to the first two sentences, the verb and a determiner. We quantified the constraint of these discourse contexts by carrying out a second cloze norming study. In this study, lists were divided into thirds to minimize time demands on any individual participant. After exclusions, between 51 and 69 participants provided completions for each context.

*(b) List composition and counterbalancing*

In the main manuscript, we report contrasts between three experimental conditions: *Expected*, *Unexpected plausible* and *Implausible*. We created 100 *expected* discourse scenarios by pairing 100 high constraint discourse contexts with the noun that was produced most frequently in the second cloze norming study. We created 100 *unexpected plausible* scenarios by pairing the same high constraint contexts with direct object nouns of low cloze values but that were still plausible in context. Finally, we created 150 *implausible* scenarios in which the noun violated the animacy constraints of the prior verb. In 100 of these *implausible* scenarios, we used the same high constraint contexts as the other two conditions. The other 50 *implausible* scenarios were created using 50 additional high constraint discourse contexts. These had served as fillers in our previous ERP study (Kuperberg, Brothers, & Wlotko, 2020), but, in the current study, we made the *a priori* decision to include them in our analyses so that we could maximize our statistical power to detect underlying neuroanatomical sources.

In addition to these 350 experimental scenarios, we also constructed an additional 350 scenarios with low constraint contexts. Of these additional low constraint scenarios, 150 were plausible and constituted a fourth *low constraint unexpected* condition (see Kuperberg et al., 2020). Although we do not report analyses involving these stimuli in the present manuscript, they were considered a fourth condition for the purpose of counterbalancing, and we plan to report analyses that include this condition in a separate manuscript. The remaining 200 low constraint scenarios were implausible and were considered fillers. These fillers were based on the *low constraint anomalous* condition that was included in our previous ERP study (Kuperberg et al., 2020).

The full set of scenarios were divided into four lists, which were rotated across participants. Each list included 200 scenarios in total: our 100 experimental scenarios in which critical words followed high constraint contexts (25 *Expected*, 25 *Unexpected plausible*, and 50 *Implausible*), and 100 additional scenarios in which critical words followed low constraint contexts (50 *Low constraint plausible* and 50 *Low constraint implausible*). Therefore, each list contained the same number of plausible and implausible critical nouns, and the same number of high constraint and low constraint discourse contexts. In creating the lists, we were able to partially counterbalance the introductory two sentences of each scenario, the verbs that preceded the critical nouns in the third sentence, and critical nouns (across the *unexpected plausible* and *implausible* conditions) so that participants saw these aspects of the scenario only once, but across all participants, they were seen in more than one condition.

2. Spatial search region used for MEG source localization

**Supplementary Figure 1: Search region**

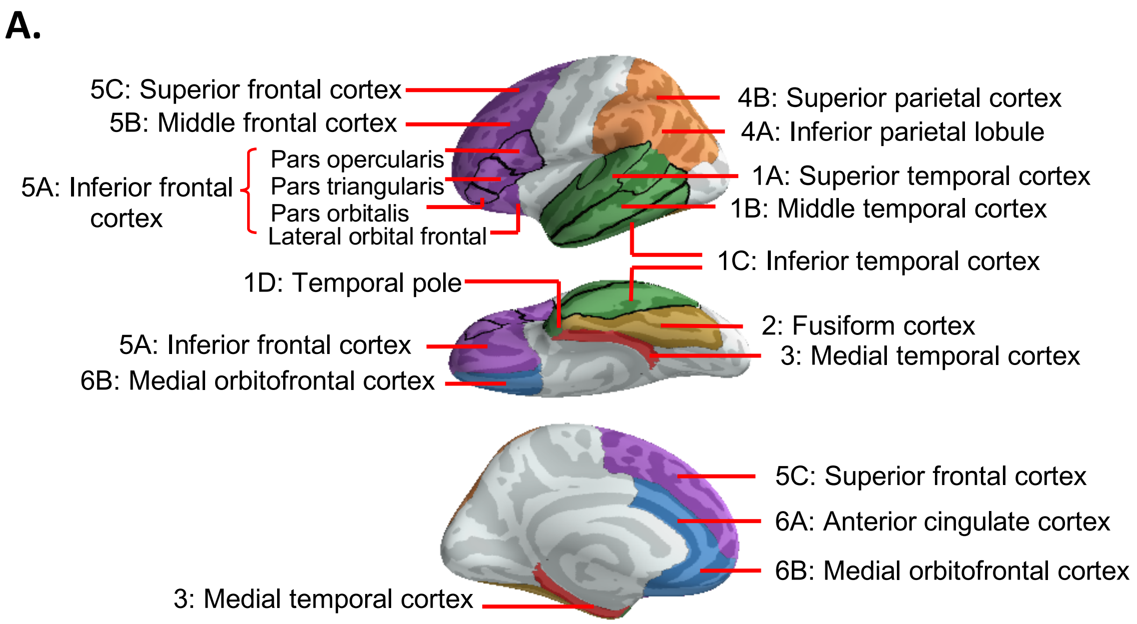


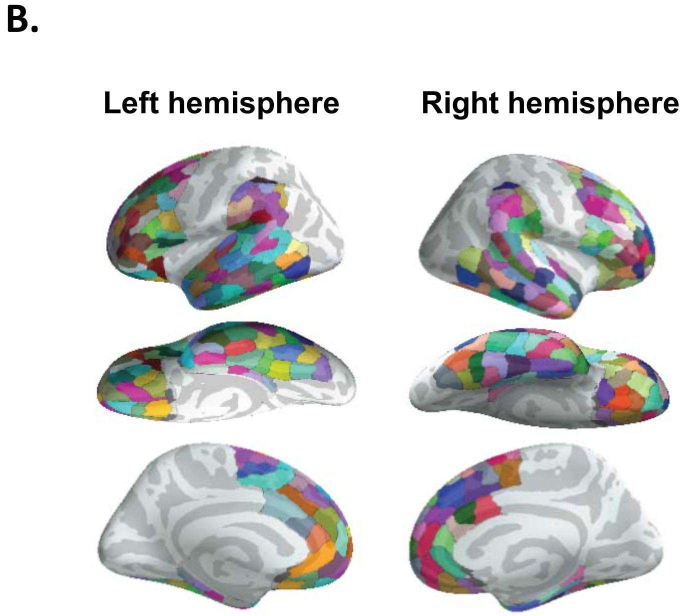


**A. Left-lateralized search region used to carry out MEG statistical analysis.** These regions were defined on the “fsaverage” FreeSurfer surface (Fischl, Sereno, & Dale, 1999) using the Desikan–Killiany atlas (Desikan et al., 2006). Supplementary Table 1 lists the correspondence between the names of the regions indicated here, and the nomenclature of the equivalent regions in the Desikan–Killiany atlas.
***B.* Patches defined on the “fsaverage” FreeSurfer surface.** These patches were used in a spatial cluster-based permutation test to account for multiple comparisons. In total, 140 patches were defined on the left hemisphere within our *a priori* search area, and 141 patches were defined on the right hemisphere for an exploratory analysis.

**Supplementary Table 1. A detailed description of the left lateralized search region used for MEG analysis.**

| Region | Number | Desikan-Killiany atlas |
|---|---|---|
| ***Lateral temporal cortex*** | | |
| #Superior temporal cortex | 1A | superiortemporal-lh<br>bankssts-lh<br>transversetemporal-lh |
| Middle temporal cortex | 1B | middletemporal-lh |
| Inferior temporal cortex | 1C | inferiortemporal-lh |
| Temporal pole | 1D | temporalpole-lh |
| ***Ventral temporal cortex*** | | |
| Fusiform cortex | 2 | fusiform-lh |
| ***Medial temporal cortex*** | | |
| Medial temporal cortex | 3 | parahippocampal-lh<br>entorhinal-lh |
| ***Lateral parietal cortex*** | | |
| Inferior parietal lobule | 4A | inferiorparietal-lh<br>supramarginal-lh |
| Superior parietal cortex | 4B | superiorparietal-lh |
| ***Lateral frontal cortex*** | | |
| ^Inferior frontal cortex | 5A | parsorbitalis-lh<br>parstriangularis-lh<br>parsopercularis-lh lateralorbitofrontal-lh<br>frontalpole-lh |
| Middle frontal cortex | 5B | caudalmiddlefrontal-lh<br>rostralmiddlefrontal-lh |
| *Superior frontal cortex | 5C | superiorfrontal-lh |
| ***Medial frontal cortex*** | | |
| **Anterior cingulate cortex | 6A | rostralanteriorcingulate-lh<br>caudalanteriorcingulate-lh |
| Medial orbitofrontal cortex | 6B | medialorbitofrontal-lh |

The names and numbers of each region correspond to those indicated in Supplementary Figure 1, which illustrates the full search region. Regions were defined on the "fsaverage" FreeSurfer surface (Fischl et al., 1999), using the Desikan–Killiany atlas (Desikan et al., 2006). This table lists the correspondences between the numbers and names of the regions shown in Supplementary Figure 1 and the names of the regions from the Desikan–Killiany atlas.

#We grouped multiple gyri defined in the Desikan-Killiany atlas into one cortical region. The gyrus includes the part visible on the pial view plus its adjacent banks of the sulci delineating this gyrus.

^The lateral portion of the orbitofrontal cortex and frontal pole are included in the left inferior frontal cortex.

*Both lateral and medial surfaces are included within the superior frontal region.

**Both anterior and middle surfaces are included within the anterior cingulate cortex.

### Supplementary Results

1. Supplementary behavioral results

On average, participants provided accurate plausibility judgments on 88.3% of trials (SD: 11.1%). There was a significant main effect of Scenario Type, $F(1,31) = 45.85$, $p < 0.001$, $eta^2 = 0.60$. Follow-up pairwise comparisons indicated that participants were most accurate in responding "Yes" to the *expected* scenarios (Mean: 95.6%; SD: 5.6%), followed by responding "No" to the *implausible* scenarios (Mean: 88.3%, SD: 8.0%), and they were least accurate in responding "Yes" to the *unexpected plausible* scenarios (Mean: 81.0%, SD: 13.0%). On average, 80.4% of the comprehension questions were answered correctly (SD: 14.2%), which suggests that participants were attending to both the introductory context and the final critical sentence.

2. Exploratory analysis of ERP and source-localized MEG data between 200-300ms

As shown in Figure 3A in the main manuscript, there appeared to be a divergence between the ERP waveforms evoked by the *expected* words and the two other conditions between 200-300ms (before the 300-500ms N400 time-window). This effect had a broad distribution that was maximal over anterior and central electrodes. An analysis in which we collapsed across all spatial regions and all time-points between 200-300ms confirmed a significant difference between the *expected* and both the *unexpected plausible* ($t_{(31)} = 3.62$, $p < .001$, $d = 1.30$) and the *implausible* ($t_{(31)} = 5.75$, $p < .001$, $d = 2.07$) words, but no difference between the *unexpected plausible* and the *implausible* words ($t_{(31)} = 0.99$, $p = 0.33$, $d = 0.36$).

This finding replicates previous ERP studies that have also observed effects of predictability before the classical 300-500ms (N400) time-window. These early effects have been interpreted as reflecting facilitated processing of predictable inputs at sublexical levels of representation. For example, the reduced negativity to *expected* words has been interpreted as a reduced N250 (Brothers, Swaab, & Traxler, 2015; Lau, Holcomb, & Kuperberg, 2013)—an ERP component that is thought to reflect sublexical orthographic processing (Grainger & Holcomb, 2009; Holcomb & Grainger, 2006; Kiyonaga, Grainger, Midgley, & Holcomb, 2007), as well as a reduced phonological mismatch negativity, reflecting facilitated phonemic processing (Connolly & Phillips, 1994; see also van den Brink, Brown, & Hagoort, 2001).

Within a predictive coding framework, facilitation at sublexical levels of linguistic representation would be attributed to the reduction of sublexical (e.g., orthographic) prediction error. On this account, in high constraint contexts, the brain not only generates top-down lexico-semantic predictions that suppress prediction error produced by expected inputs at the lexico-semantic level between 300-500ms, but it also generates predictions at sublexical levels of representation that suppress prediction error produced by expected inputs at these lower levels of linguistic representation between 200-300ms.

However, there are other interpretations of these early ERP effects of predictability (see Nieuwland, 2019 for a review). For example, it is possible that, between 200-300ms, the reduced negativity to *expected* incoming words reflects an early divergence of the N400 component itself, indexing facilitated lexico-semantic processing (see Lau, Holcomb, et al., 2013 for discussion). In addition, others have argued that, instead of reflecting a reduced negativity to expected words, these early effects may, in fact, reflect an enhanced positivity to words that *confirm* prior predictions, e.g. an enhanced P2 due to a top-down attentionally-mediated extraction of visual features (Federmeier, Mai, & Kutas, 2005), or a P3b that reflects the categorization of a confirmed prediction (Molinaro & Carreiras, 2010; Roehm, Bornkessel-Schlesewsky, Rosler, & Schlesewsky, 2007; Vespignani, Canal, Molinaro, Fonda, & Cacciari, 2010).

As shown in Figure 5 (left), the MEG source-localized dynamic Statistical Parametric Maps (dSPMs) within this 200-300ms time-window reveal effects that appear to be more compatible with a reduction (rather than an enhancement) of neural activity to the *expected* words. However, they do not shed light on whether these reductions in activity reflect facilitation at the lexico-semantic or sublexical levels of representation. The *expected* words appear to show less activity than both the *unexpected plausible* and *implausible* words in superior temporal, medial temporal and posterior (occipitotemporal) fusiform cortices. However, paired statistical contrasts revealed significant effects only within superior and medial temporal cortices, and only when comparing the *implausible* and *expected* inputs. The failure to find statistical effects in other regions and/or other contrasts may be due to a lack of power: the ERP effect between 200-300ms is much smaller than the later N400 effect. Therefore, in order to accurately source-localize this early effect, it will be important to carry out an MEG study that has a very large number of items per condition, and to fully counterbalance the same critical words across levels of predictability.

Such a study would provide important data for resolving controversies regarding the validity and functional interpretation of early predictability effects.

3. MEG source-localization videos

In the attached videos, we show averaged dSPM source activations from 0-1000ms, in 10ms bins, after critical word onset in each of the three experimental conditions.

4. Exploratory analysis of a subset of the MEG data

As noted in the main manuscript, the critical words in the *expected* scenarios were more frequent and had smaller orthographic neighborhoods than the critical words in the two other conditions. These lexical differences between the *expected* words and the other conditions were a function of our counterbalancing scheme, which required us to use the same high constraint discourse contexts across conditions. In addition, the *implausible* condition included twice as many trials as the two other conditions (50 *versus* 25). This ensured that each participant saw an equal proportion of plausible and implausible scenarios. We made the *a priori* decision to include all 50 implausible scenarios in our main analyses in order to maximize power.

We think that it is unlikely that either of these factors systematically influenced our results. First, although previous ERP studies have shown that both frequency and orthographic neighborhood can modulate the N400 (Barber, Vergara, & Carreiras, 2004; Holcomb, Grainger, & O'Rourke, 2002; Laszlo & Federmeier, 2011, 2014; Rugg, 1990; Young & Rugg, 1992), these lexical effects tend to be much smaller than the effects of predictability and contextual plausibility on the N400. Second, we used *non-parametric* mass univariate statistical tests to analyze our data, which are robust to differences in numbers of trials between conditions (the null distribution was created by randomly permutating the same dataset with the same signal-to-noise ratio).

Nonetheless, to alleviate any concerns that either unmatched lexical properties or unequal trial numbers influenced our results, we conducted an additional analysis. In this analysis, within each participant, we randomly selected a subset of 25 *implausible* scenarios with critical words that matched the *expected* critical words on lexical characteristics (i.e. word length, word frequency and orthographic neighborhood size). We then computed the ERFs of these randomly selected *implausible* trials within each participant, and compared the source-localized activity within the 300-500ms time-window with activity produced by the *expected* trials across all participants. This analysis revealed the same pattern of results as that reported in the main manuscript.

5. Exploratory analysis of MEG source-level activity over the right hemisphere

**Supplementary Figure 2. Exploratory analysis of MEG source-level activity over the right hemisphere produced by the unexpected plausible and the expected critical words.**

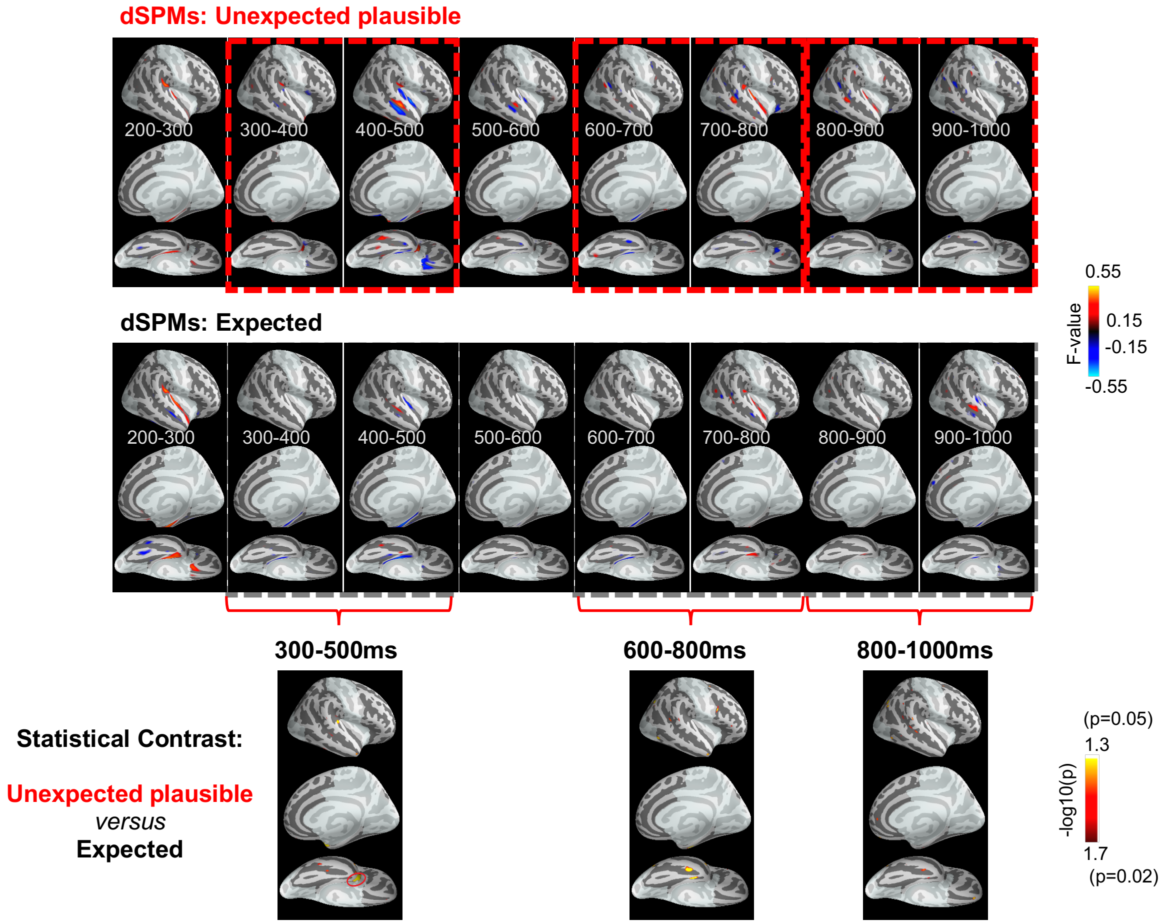


*Top and middle:* Signed dynamic Statistical Parametric Maps (dSPMs) produced by the *unexpected plausible* and the *expected* critical words, shown at 100ms intervals from 200ms until 1000ms. All dSPMs are thresholded at 0.15, with red indicating outgoing currents and blue indicating ingoing currents. *Bottom:* Statistical maps contrasting the *unexpected plausible* and *expected* critical words within our three time-windows of interest: 300-500ms, 600-800ms, and 800-1000ms. Red circles indicate regions that reached cluster-level significance. Within the 300-500ms time window, the *unexpected plausible* critical words evoked significantly more activity than the *expected* critical words within the right temporal pole. No significant effects were found within the 600-1000ms time-window.

**Supplementary Figure 3. Exploratory analysis of MEG source-level activity over the right hemisphere produced by the implausible and the expected critical words.**

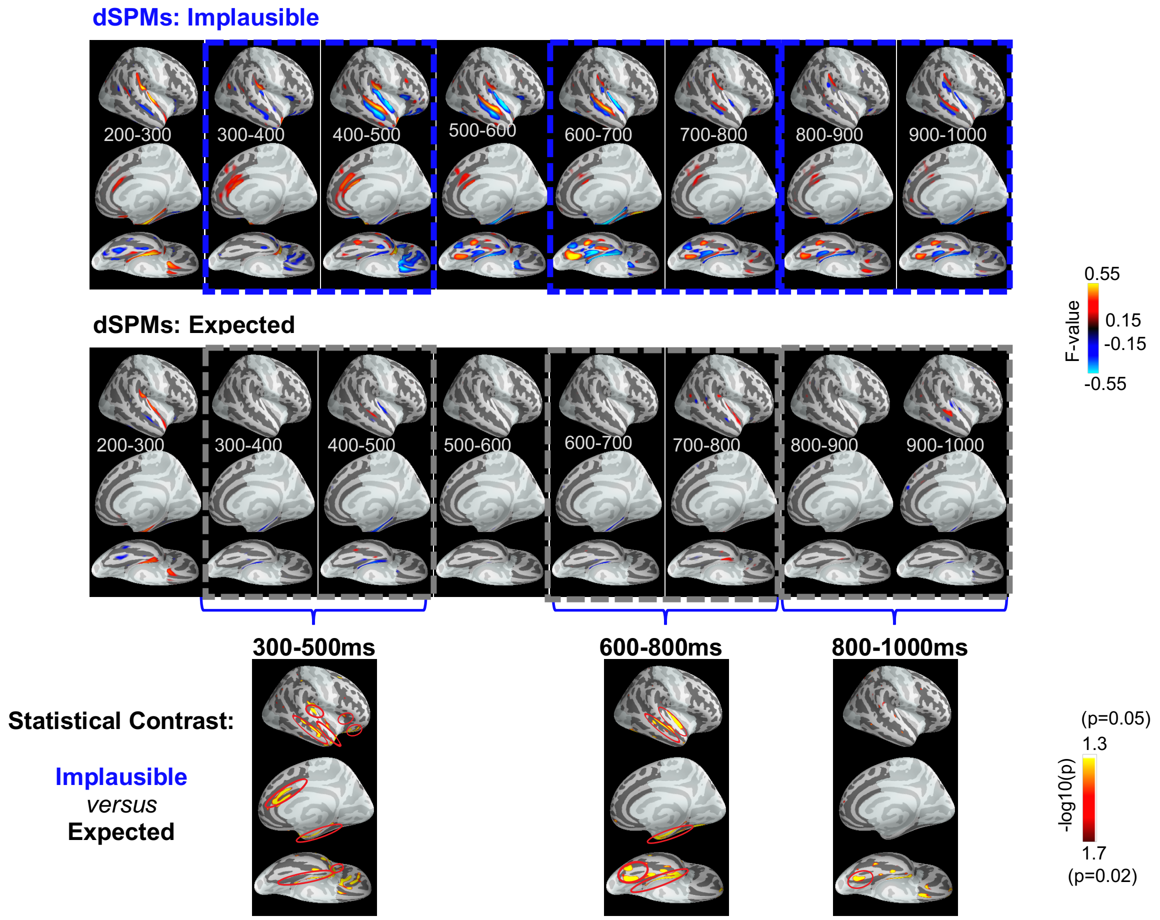


*Top and middle:* Signed dynamic Statistical Parametric Maps (dSPMs) produced by the *implausible* and the *expected* critical words, shown at 100ms intervals from 200ms until 1000ms. All dSPMs are thresholded at 0.15, with red indicating outgoing currents and blue indicating ingoing currents. *Bottom:* Statistical maps contrasting the *implausible* and *expected* critical words within our three time-windows of interest: 300-500ms, 600-800ms, and 800-1000ms. Red circles indicate regions that reached cluster-level significance. Within the 300-500ms time window, the *implausible* critical words evoked significantly more activity than the *expected* critical words within the right lateral temporal cortex, the right anterior inferior frontal cortex, and the right anterior cingulate cortex. This contrast also revealed an effect in the right medial temporal cortex, which was driven by dipoles going in opposite directions to the *implausible* (outgoing) and the *expected* (ingoing) critical words. The locations of these effects were similar to those observed over the left hemisphere, but they appeared to be less robust. Between 600-1000ms, the *implausible* critical words produced more activity within the right posterior fusiform cortex than the *expected* critical words (significant in both the 600-800ms and the 800-1000ms windows). Between 600-800ms, the *implausible* critical words also produced significantly more activity than the *expected* critical words within the right superior temporal and medial temporal cortices.

**Supplementary Figure 4. Exploratory analysis of MEG source-level activity over the right hemisphere produced by the unexpected plausible and the implausible critical words.**

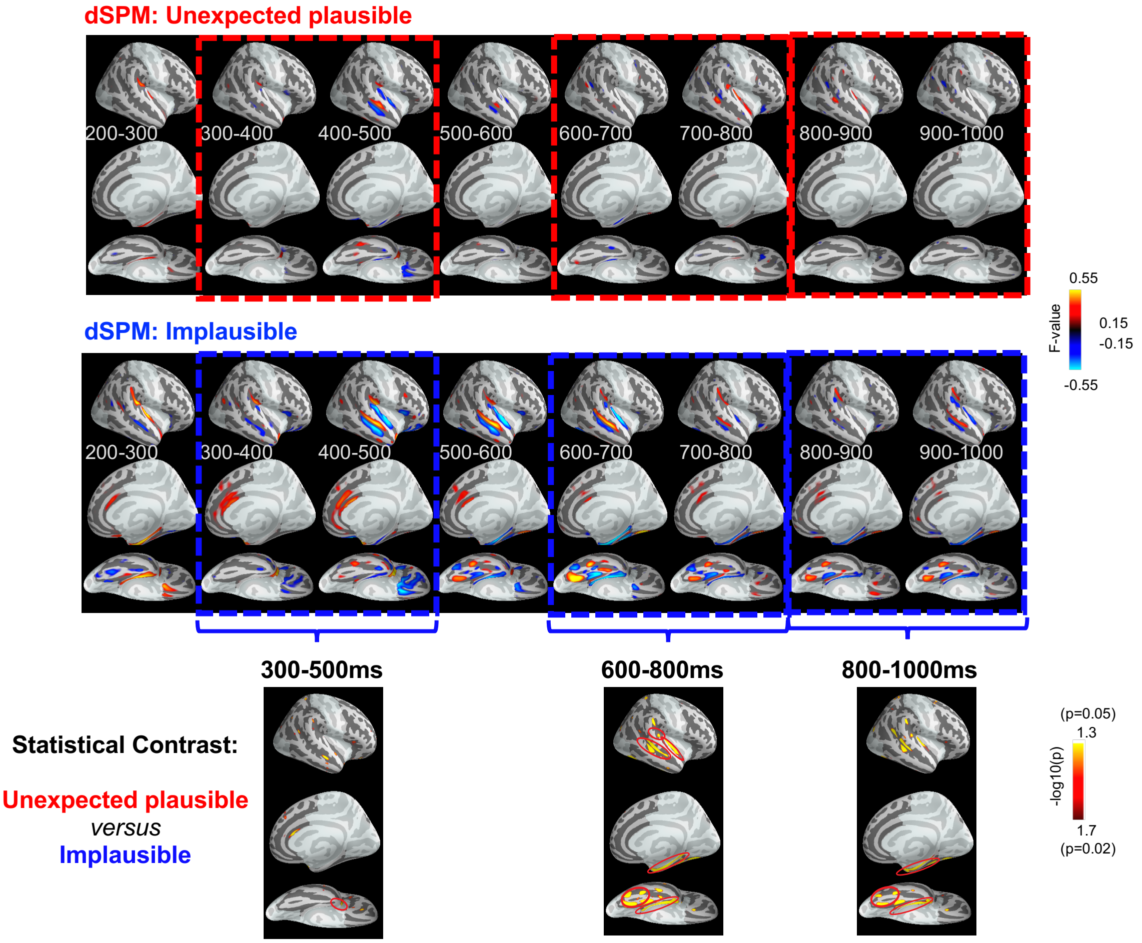


*Top and middle:* Signed dynamic Statistical Parametric Maps (dSPMs) produced by the *unexpected plausible* and *implausible* critical words, shown at 100ms intervals from 200ms until 1000ms. All dSPMs are thresholded at 0.15, with red indicating outgoing currents and blue indicating ingoing currents. *Bottom:* Statistical maps contrasting the *unexpected plausible* and *implausible* critical words within our three *a priori* time windows of interest: 300-500ms, 600-800ms, and 800-1000ms. Red circles indicate regions that reached cluster-level significance. Within the 300-500ms time window, the *implausible* critical words evoked significantly more activity than the *unexpected plausible* critical words within the right temporal pole. Between 600-1000ms, the *implausible* critical words produced more activity within the right posterior fusiform and medial temporal regions than the *unexpected plausible* critical words (significant in both the 600-800ms and the 800-1000ms windows). Between 600-800ms, *implausible* critical words also produced significantly more activity within the right superior temporal cortex than the *unexpected plausible* critical words.

### Supplementary Discussion

1. The relationship between the MEG and ERP findings

As noted in the main manuscript, although MEG and ERP both measure phase-locked evoked activity, they do not capture precisely the same underlying signal, and so they do not always pattern together (Ahlfors, Han, Belliveau, & Hämäläinen, 2010). By simultaneously collecting ERP and MEG data using the same stimuli in the same participants, we were able to directly compare the ERP and MEG effects. In general, the two methods revealed similar patterns of modulation across the three conditions (e.g. graded increases of activity within the N400 time-window, and spatially distinct effects to the *unexpected plausible* and *implausible* continuations in the late time-window). However, there were also some interesting differences.

First, the effect of implausibility (*implausible* versus *unexpected plausible*) on the ERP N400 (shown in Figure 1, main manuscript) appeared to be much smaller than on the the sensor-level MEG N400 (shown in Figure 2, main manuscript). We suggest that this is because the N400 ERP component evoked by the *implausible* critical words was artificially reduced at the scalp surface as a result of spatiotemporal overlap with the subsequent *late posterior positivity/P600* ERP component that was produced by these continuations. The N400 and *late posterior positivity/P600* ERP components both have posterior scalp distributions. However, because they have opposite polarities, they can cancel each other out at the scalp surface (Brouwer & Crocker, 2017; Kuperberg, Kreher, Sitnikova, Caplan, & Holcomb, 2007). This type of "component overlap" is less of an issue for MEG for two reasons. First, the signal detected by gradiometer MEG sensors does not carry information about the polarity of the underlying dipoles; that is, sensor-level evoked MEG responses reflect the overall magnitude of activity, regardless of the direction of the underlying currents. Therefore, unlike ERP responses, there is no cancellation of the MEG signal at the scalp surface. Second, MEG has a better spatial resolution than EEG because magnetic fields are less distorted than electric fields by the conductivities of the skull and scalp. Therefore, evoked MEG responses that originate from spatially distinct underlying sources are less likely than ERP responses to overlap spatially at the scalp surface within the same time-window.

Note that this account of ERP component overlap implies that the *late posterior positivity/P600* ERP produced by the *implausible* words began within the N400 time window. This, in turn, implies that the conflict that triggered the *late posterior positivity/P600* effect was detected in the 300-500 (N400) time window. This early detection of conflict may have been indexed by the anterior cingulate response to the *implausible* (versus *expected*) words, detected by MEG between 300-500ms.

A second difference between the MEG and ERP findings was in the magnitude of the late effects observed between 600-1000ms. In ERPs, the magnitude of these late effects (both the *late frontal positivity* evoked by the *unexpected plausible* continuations and the *late posterior positivity/P600* evoked by the highly *implausible* continuations) were generally larger than the MEG effects for the same contrasts observed within the same late time window. The relative insensitivity of MEG to neural effects that manifest in the ERP waveform as robust positive-going components has been noted before (Ahlfors, Han, Lin, et al., 2010). For example, MEG is relatively insensitive to the well-known domain-general P3b effect (Siedenberg, Goodin, Aminoff, Rowley, & Roberts, 1996), to which the late posterior positivity/P600 is thought to be functionally related (Coulson, King, & Kutas, 1998; Osterhout, Kim, & Kuperberg, 2012; Sassenhagen & Fiebach, 2019; Sassenhagen, Schlesewsky, & Bornkessel-Schlesewsky, 2014). One possible reason for this is that, unlike ERPs, which index activity originating from both sulci and gyri, MEG is insensitive to radial sources from gyri (Ahlfors, Han, Belliveau, et al., 2010). In addition, in MEG, tangential sources on opposing sides of sulci often cancel out (Ahlfors, Han, Lin, et al., 2010). For both these reasons, MEG is relatively insensitive to activity that stems from *extended* regions of cortex that cut across multiple gyri and sulci, and that may produce large late positivity ERP effects.

These differences between ERP and MEG may also have functional implications; that is, the two measures may not necessarily index precisely the same *cognitive* mechanisms. For example, consider the process of reanalysis that is thought to be triggered by highly *implausible* inputs between 600-1000ms (Brothers, Zeitlin, Choi Perrachione, Choi, & Kuperberg, in press; van de Meerendonk, Kolk, Chwilla, & Vissers, 2009). As recently discussed (Alexander, Brothers, & Kuperberg, Submitted), reanalysis is likely to comprise at least two closely-linked but distinct sets of mechanisms: (a) sequential decision-making and active information sampling (Gottlieb, 2012; Gottlieb & Oudeyer, 2018), which involve the accumulation of probabilistic evidence for a linguistic error (conflict with the structure and state of the generative model), and the reallocation of attention when this evidence crosses a particular threshold, and (b) linguistic reprocessing of the input itself. These processes are likely to proceed in parallel and to interact closely with one another. The first set of mechanisms has been linked to the P3b (Kelly & O'Connell, 2013; O'Connell, Dockree, & Kelly, 2012; Twomey, Murphy, Kelly, & O'Connell, 2015) as well as the “error positivity” (Desender, Murphy, Boldt, Verguts, & Yeung, 2019; Murphy, Robertson, Harty, & O'Connell, 2015). Evidence accumulation is also closely linked to the phasic release of norepinephrine from the locus coeruleus (Dayan & Yu, 2006), which is also linked to the P3b (de Gee, Correa, Weaver, Donner, & van Gaal, 2021; Nieuwenhuis, Aston-Jones, & Cohen, 2005), and may function to increase the gain of processing within widespread regions of cortex (Aston-Jones & Cohen, 2005; Sara, 2009). It is therefore possible that ERP recordings in this late time-window (i.e. the late positivities) were more sensitive to these widespread decision-making and attention mechanisms, while the MEG evoked response within the posterior fusiform cortex within the same time window was relatively more sensitive to the more localized process of orthographically reanalyzing the input.

<u>2. The retention of dipole polarity when carrying out MEG source localization analyses</u>

When carrying out distributed source localization of our MEG data, we chose to retain the polarity (signed values) of the estimated dipoles. This allowed us not only to estimate the magnitude of activity across experimental conditions, but also to determine the direction of the underlying dipoles, i.e. whether the current was ingoing or outgoing, relative to the cortical surface. At a neurophysiological level, the polarity of a dipole is determined both by the precise configuration of the pyramidal cells and the direction of the intracellular currents they generate (Lopes da Silva, 2010). Although the precise mechanisms that give rise to differences in dipole polarity are unclear, it is likely that systematic *differences* in dipole polarity between conditions and/or time-windows has some functional significance. As we discussed next, our results revealed some interesting patterns.

*(a) Different conditions can produce dipoles of opposite polarities within the same time-windows and neuroanatomical regions*

In several cases, we found that the statistical differences between two conditions within a given region and time-window were driven by dipoles going in opposite directions to each condition. For example, in the N400 time-window, the effect in the left medial temporal cortex, was driven not only by a dipole to the unpredictable words (both to the *unexpected plausible* and *implausible* words), but also by a dipole to the *expected* words. As discussed in the main manuscript, we speculated that the dipole produced by the *expected* words indexed the detection of a match (so-called "resonance"; Carpenter & Grossberg, 1993) between pre-activated activity within this medial temporal region, and the expected bottom-up input.

Beyond its theoretical implications, our finding that effects can be driven by dipoles going in opposite directions also has methodological implications. Previous intracranial studies have also reported effects within medial temporal cortex within the N400 time-window, and it has been noted that different electrodes within these medial temporal regions produce local field potentials of opposite polarities to incongruous and expected words (McCarthy, Nobre, Bentin, & Spencer, 1995). However, previous MEG studies using distributed source localization have failed to report effects within the medial temporal cortex within the N400 time window, either in semantic priming paradigms (Lau, Gramfort, Hämäläinen, & Kuperberg, 2013; Lau, Weber, Gramfort, Hämäläinen, & Kuperberg, 2016) or during sentence comprehension (Maess, Herrmann, Hahne, Nakamura, & Friederici, 2006). We suggest that this may because these previous MEG studies used unsigned rather than signed dipole values for source localization. The absolute values of two dipoles going in opposite directions would have therefore canceled out, and failed to show any significant difference at all to predictable versus unpredictable words.

In addition to the medial temporal effects between 300-500ms, we also observed effects that were driven by dipoles going in opposite directions across conditions within the later 600-1000ms time-window: (1) the effects within left inferior frontal and middle temporal cortices in contrasting the *unexpected plausible* and *expected* words, and (2) the effect within left inferior frontal cortex in contrasting the *implausible* and *expected* words. Once again, these differences would have been overlooked if we had only examined the absolute amplitude of the source activity.

*(b) The same condition can produce dipoles within the same region that reverse in polarity across time-windows*

In several other cases, a dipole produced by a particular condition showed a polarity reversal between the 300-500ms and the later 600-1000ms time-window. This was the case for the evoked responses produced by the *unexpected plausible* words within the left lateral temporal cortex, as well as the response produced by the *implausible* words within the left inferior frontal and medial temporal cortices. Again, although the precise mechanisms underlying these dipole reversals are unclear, they are again likely to have some functional significance. In the main manuscript, we noted that these dipole reversals were consistent with a functional distinction between early feedforward activity within the 300-500ms time-window and later feedback activity within the 600-1000ms time-window.

*(c) Methodological and theoretical implications*

Taken together, these observations add to a growing body of work suggesting that there are several advantages of retaining dipole polarity when estimating underlying neural sources. Previous MEG work that has systematically compared analyses of the same datasets using signed and unsigned estimates, has shown that retaining signed summary statistics is robust to factors such as individual neuroanatomical variability and spatial smoothing (Henson et al., 2007). In addition, a recent study showed that retaining dipole direction information allowed for a better characterization of the time course of visual word processing (Gwilliams, Lewis, & Marantz, 2016). As discussed in the Methods section of the main manuscript, the retention of dipole information also has the advantage of allowing analyses between conditions with unequal numbers of trials, in contrast to traditional methods that square positive and negative values to yield positively-signed estimate of dipole magnitude, thereby artificially inflating noise estimates in conditions with more trials. Finally, as discussed above, retaining dipole polarity ensures that effects that are driven by dipoles going in opposite directions are not cancelled out.

The present set of findings suggest that, beyond these methodological implications, a systematic examination of dipole polarity in MEG source localization may also have functional implications, yielding new insights into the cognitive mechanisms that support language comprehension.
